# Discovery of Cyclic Peptide Ligands to the SARS-CoV-2 Spike Protein using mRNA Display

**DOI:** 10.1101/2020.12.22.424069

**Authors:** Alexander Norman, Charlotte Franck, Mary Christie, Paige M. E. Hawkins, Karishma Patel, Anneliese S. Ashhurst, Anupriya Aggarwal, Jason K. K. Low, Rezwan Siddiquee, Caroline L. Ashley, Megan Steain, James A. Triccas, Stuart Turville, Joel P. Mackay, Toby Passioura, Richard J. Payne

## Abstract

The COVID-19 pandemic, caused by SARS-CoV-2, has led to substantial morbidity, mortality and disruption globally. Cellular entry of SARS-CoV-2 is mediated by the viral spike protein and affinity ligands to this surface protein have the potential for applications as antivirals and diagnostic reagents. Here, we describe the affinity selection of cyclic peptide ligands to the SARS-CoV-2 spike protein receptor binding domain (RBD) from three distinct libraries (in excess of a trillion molecules each) by mRNA display. We identified six high affinity molecules with dissociation constants (*K*_D_) in the nanomolar range (15-550 nM) to the RBD. The highest affinity ligand could be used as an affinity reagent to detect spike protein in solution by ELISA, and the co-crystal structure of this molecule bound to the RBD demonstrated that it binds to a cryptic binding site, displacing a β-strand near the C-terminus. Our findings provide key mechanistic insight into the binding of peptide ligands to the SARS-CoV-2 spike RBD and the ligands discovered in this work may find future use as reagents for diagnostic applications.

## Introduction

The COVID-19 pandemic, caused by infection with the severe acute respiratory syndrome related coronavirus-2 (SARS-CoV-2), has led to widespread morbidity and mortality and has resulted in crippling effects on the global economy. At the time of writing there had been > 77 million confirmed cases of COVID-19 and >1.7 million deaths resulting from SARS-CoV-2 infections. Since the first reported case of COVID-19 in the province of Hubei, China in December 2019, there has been an intense global research effort centered on the development of an effective vaccine for the control of the disease. Very recently, two mRNA vaccines (BNT162b2 and mRNA-1273, developed by Pfizer/BioNTech and Moderna, respectively) and a chimpanzee adenovirus-vectored vaccine (ChAdOx1 nCoV-19, from Oxford University and AstraZeneca) have completed phase III clinical trials and demonstrated a favorable safety profile and promising efficacy.^1–3^ In addition to the requirement for a vaccine, effective antiviral drugs are also critically needed for COVID-19 treatment; these drugs will be particularly important for use in unvaccinated individuals, in cases where the efficacy of a vaccine wanes or the virus develops resistance to vaccine-induced responses.^4^

While several candidate antiviral molecules are already in clinical trials, none were developed specifically for treatment of SARS-CoV-2 infection. Moreover, even the most promising of these, including remdesivir, hydroxychloroquine, lopinavir-ritonavir and type I interferon therapy, have been deemed ineffective or only modestly efficacious in clinical trials.^5–13^ Thus, in addition to investigations into the repurposing of existing drugs, there is an urgent need for the development of novel molecules that specifically target SARS-CoV-2 for the treatment of COVID-19. These molecules may also find use for immediate deployment in situations where SARS-CoV-2 becomes endemic, including for pre-exposure prophylaxis in high-risk populations, or for further SARS-CoV outbreaks, e.g. through future zoonoses.

The critical initiating step in viral infection involves entry of SARS-CoV-2 into human cells, a process that is mediated by interaction between the receptor binding domain (RBD) of the viral spike protein with cell-surface angiotensin converting enzyme 2 (ACE2) (Figure 1).^14^ Crystal structures of RBD-ACE2 complexes have elucidated the mechanism of this interaction and shown that the RBD-binding site on ACE2 is located on an N-terminal helical region, distinct from the catalytic site (Figure 1).^15, 16^ Importantly, it has recently been demonstrated that blocking this interaction (through addition of either soluble RBD protein or soluble ACE2 protein) potently inhibits viral infection of cultured cells in a dose dependent manner.^17–19^ These data provided early evidence that the RBD-ACE2 interaction was a valid target for the development of novel antivirals against SARS-CoV-2 infection. Several approaches have since been adopted for the discovery of novel RBD binding ligands, including full antibodies,^20–28^ as well as antibody fragments such as VH domains,^29, 30^ nanobodies^31–35^ and monobodies.^36^ In addition, engineered ACE2 mimetics/decoy receptors,^37–39^ *de novo* designed miniproteins,^40, 41^ peptide-based ACE2 mimetics^42, 43^ and synthetic peptide libraries^44^ have also been reported that are capable of binding the SARS-CoV-2 spike RBD.

**Figure 1.**
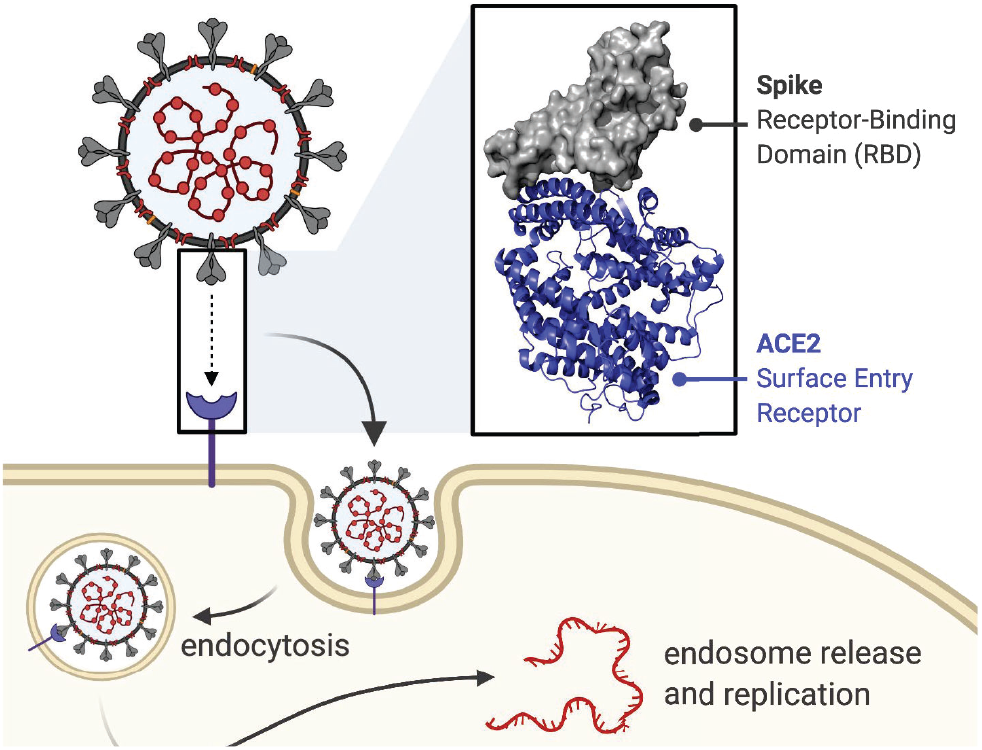
Cartoon depicting the interaction between the SARS-CoV-2 trimeric spike protein and the human ACE2 receptor, a key step in viral entry (endocytic entry route shown). Inset: crystal structure (PDB: 6LZG) showing the interaction between the N-terminal helix of ACE2 (blue) and the SARS-CoV-2 spike RBD (grey).

Macrocyclic peptides are a class of molecules demonstrated to be highly effective at disrupting protein-protein interactions, particularly in cases such as the spike-ACE2 interaction where a defined binding pocket is lacking.^45–54^ In this work we explored this chemotype for the development of SARS-CoV-2 RBD-binding molecules that block the spike-ACE2 interaction, with a view to discovering novel inhibitors of viral entry. To discover novel cyclic peptides we employed cyclic peptide mRNA display, an approach that enables the generation of libraries of >10^12^ macrocyclic peptides that can be subsequently selected against the target of interest, in our case the RBD of the spike protein of SARS-CoV-2 (Figure 1).

## Results and Discussion

To identify ligands to the SARS-CoV-2 RBD, we performed three parallel affinity selections using very high diversity macrocyclic peptide libraries (Figure 2A). Two of these were genetically reprogrammed <u>Ra</u>ndom non-standard <u>P</u>eptide <u>I</u>ntegrated <u>D</u>iscovery (RaPID) libraries, comprising thioether-closed macrocyclic peptides (one initiated with *N-*chloroacetyl-L-tyrosine and one initiated with *N-*chloroacetyl-D-tyrosine). The third comprised disulfide-closed macrocyclic peptides. In each case, a semi-random DNA library was transcribed into mRNA, followed by covalent ligation to puromycin and *in vitro* translation to yield a cyclic peptide-mRNA fusion library in excess of 10^12^ unique molecules. Following counter selection (to remove streptavidin ligands), each library was panned against biotinylated SARS-CoV-2 RBD immobilized on streptavidin beads, and an enriched DNA library was recovered by RT-PCR. After seven iterative rounds of this process, the final DNA library was sequenced to identify peptide ligands predicted to bind to SARS-CoV-2 RBD with high affinity (see Supporting Information). From this sequencing we chose nine diverse and enriched peptides for further evaluation: three L-tyrosine initiated, three D-tyrosine initiated, and three disulfide closed cyclic peptides (Figure 2B).

**Figure 2.**
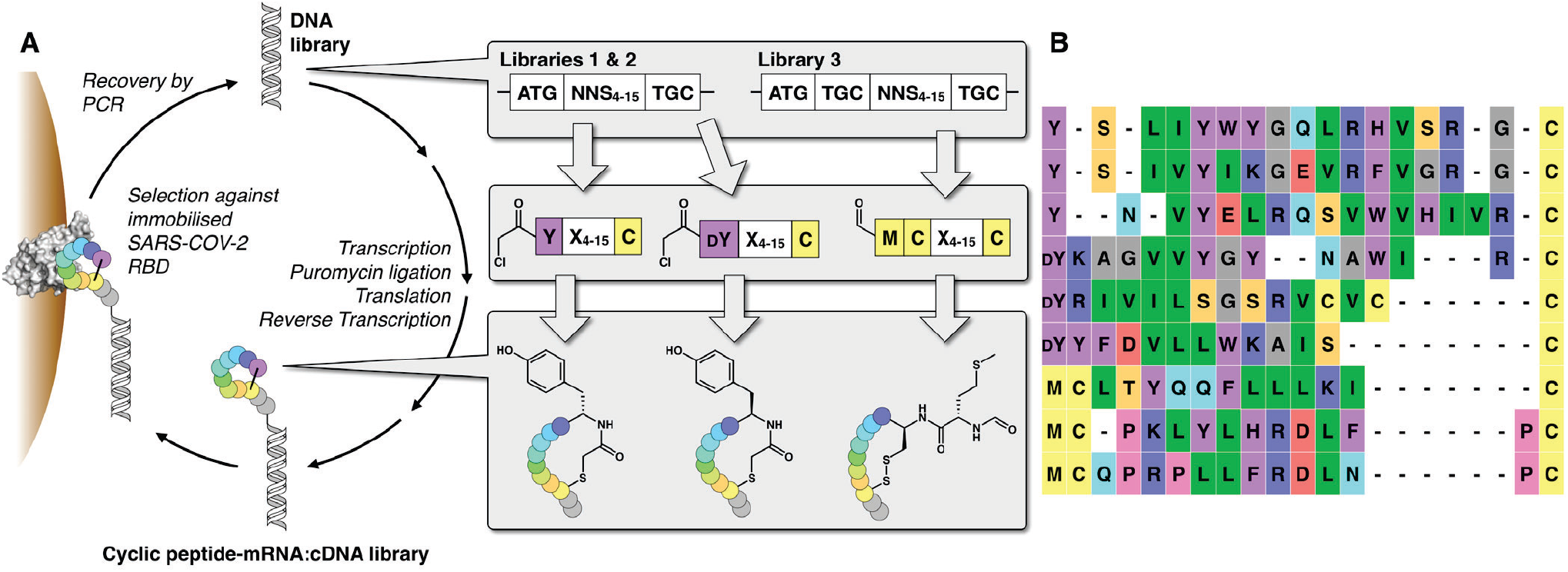
A) Schematic of the cyclic peptide mRNA display technology used. DNA libraries incorporating 4-15 randomized NNS (N = A, C, G, or T; S = C or G) codons were transcribed into mRNA, covalently ligated to puromycin (to allow conjugation between each mRNA and its cognate peptide), translated in *in vitro* reactions and reverse transcribed to afford very high diversity (>1012 molecules) peptide-mRNA:cDNA libraries. Iterative rounds of affinity selection against recombinant SARS-CoV-2 RBD protein followed by recovery of the DNA by PCR and re-synthesis of the peptide-mRNA:cDNA library were conducted to enrich for SARS-COV-2 RBD ligands. In two libraries, the initiating *N*-formyl-methionine residue was genetically reprogrammed to *N*-chloroacetyl-L-tyrosine or *N*-chloroacetyl-D-tyrosine (DY), which spontaneously cyclizes to a downstream cysteine residue to form a thioether. A third library included an additional cysteine residue affording cyclic peptides through disulfide formation. **B)** Sequence alignment of the nine enriched unique peptide sequences from each library chosen for further characterization.

The nine target cyclic peptides **1-9** were subsequently synthesized by solid-phase peptide synthesis (SPPS). Specifically, the target peptide sequences were first assembled on Rink amide resin using Fmoc-strategy SPPS (Scheme 1). For peptides **1-6** the N-termini were derivatized with chloroacetic acid (Scheme 1A), while the N-terminal methionine was *N*-acetylated in **7-9** (Scheme 1B). Each of the peptides was subsequently cleaved from resin with concomitant side chain deprotection by treatment with an acidic cocktail. It should be noted that, despite significant optimization of the solid-phase synthesis, the precursor linear peptides to **5** and **7** were generated with significant sequence truncations (as judged by LC-MS analysis after the cleavage step); these peptides were also poorly soluble in both aqueous media and organic solvents. We therefore chose to re-synthesize these two sequences with a hexaethyleneglycol solubility tag on the C-terminus. Given that the RaPID peptides were panned on the RBD bearing a large mRNA tag on the C-terminus, we were confident that this modification would not influence the binding affinity to the RBD^55, 56^ (Scheme 1). For the thioether-linked peptides **1-6**, the linear peptide precursors were cyclized by treatment with Hünig’s base in DMSO or acetonitrile/water mixtures (depending on the solubility of the linear peptides, see Supporting Information). In contrast, the disulfide-linked cyclic peptides **7-9** were generated through oxidation of the linear cleaved peptides by incubating in aqueous ammonium bicarbonate. Purification of each of the macrocyclic peptide targets by reverse-phase HPLC afforded **1-9** in 2-14% yield over the iterative SPPS and cyclization steps.

**Scheme 1.**
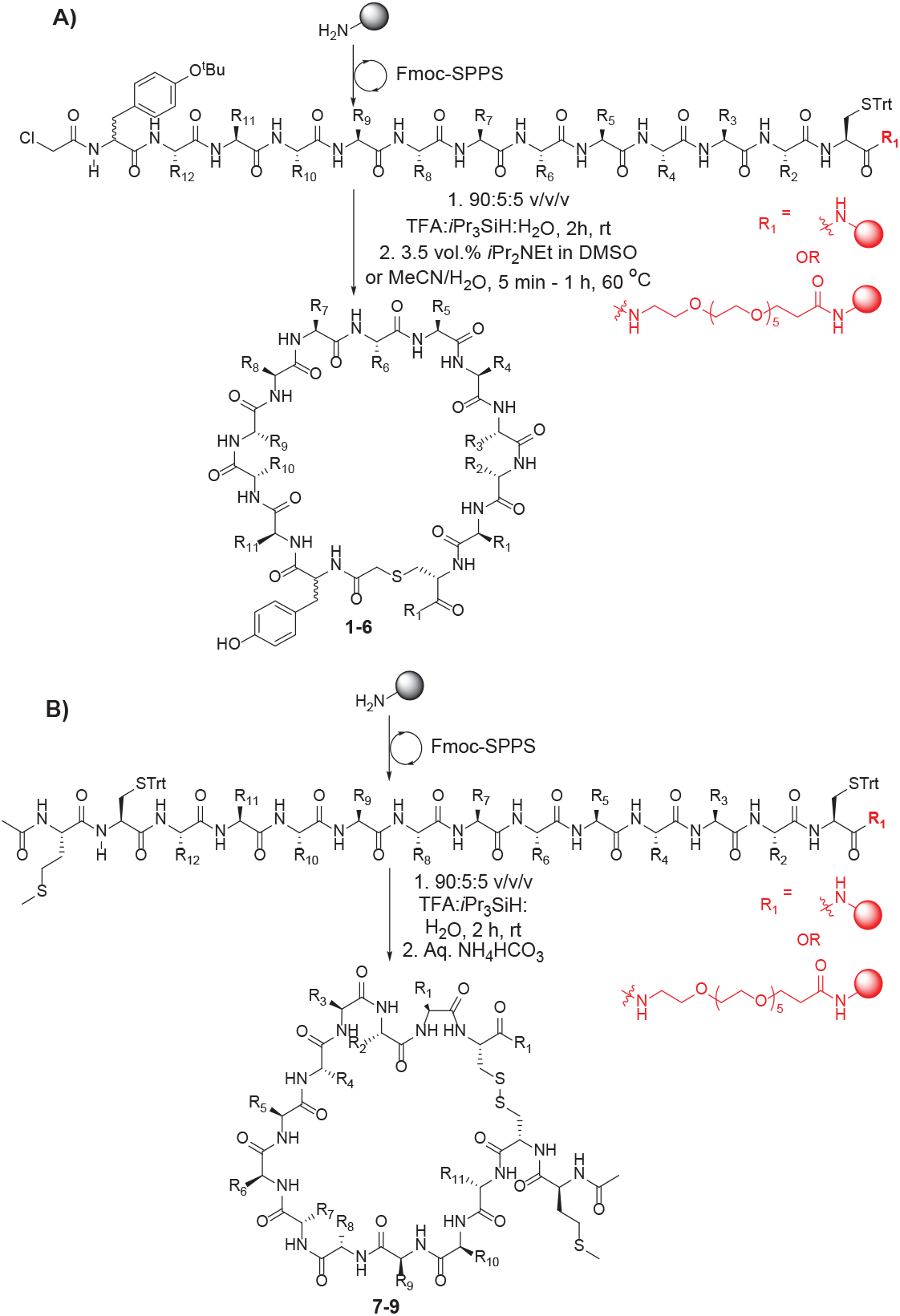
General scheme depicting the synthetic route to **A)** thioether cyclic peptide targets **1-6** and **B)** disulfide linked cyclic peptide targets **7-9**.

With the synthetic cyclic peptides **1-9** in hand, we next assessed the ability of each to bind to the RBD of SARS-CoV-2 spike protein using surface plasmon resonance (SPR). Briefly, RBD was expressed and purified before immobilizing onto a CM5 SPR chip using 1-ethyl-3-(3-dimethylaminopropyl)carbodiimide (EDC) and *N*-hydroxysuccinimide (NHS) chemistry according to the manufacturer’s protocol (see Supporting Information). Of the nine cyclic peptides screened, the six *N*-chloroacetylated peptides all exhibited high affinity for the RBD with dissociation constants (*K*_D_) in the nanomolar range (*K*_D_ = 12 – 550 nM, Table 1, see Supporting Information for SPR sensorgrams). The highest affinity of these was peptide **4**, with an observed *K*_D_ = 15 nM against RBD and a markedly low rate of dissociation (*k_off_* = 1.4 × 10^−3^ s^−1^). Interestingly, this molecule is a higher affinity cyclic analogue of a linear RBD ligand recently identified through an affinity selection-mass spectrometry approach.^44^ Peptide **5** was also notable in the series with a *K*_D_ = 76 nM against the RBD. The disulfide-closed peptides showed very high apparent RBD affinity, but also strong non-specific binding (data not shown). Based on these data we selected peptides **4** and **5** for assessment of antiviral activity against SARS-CoV-2.

**Table 1.**
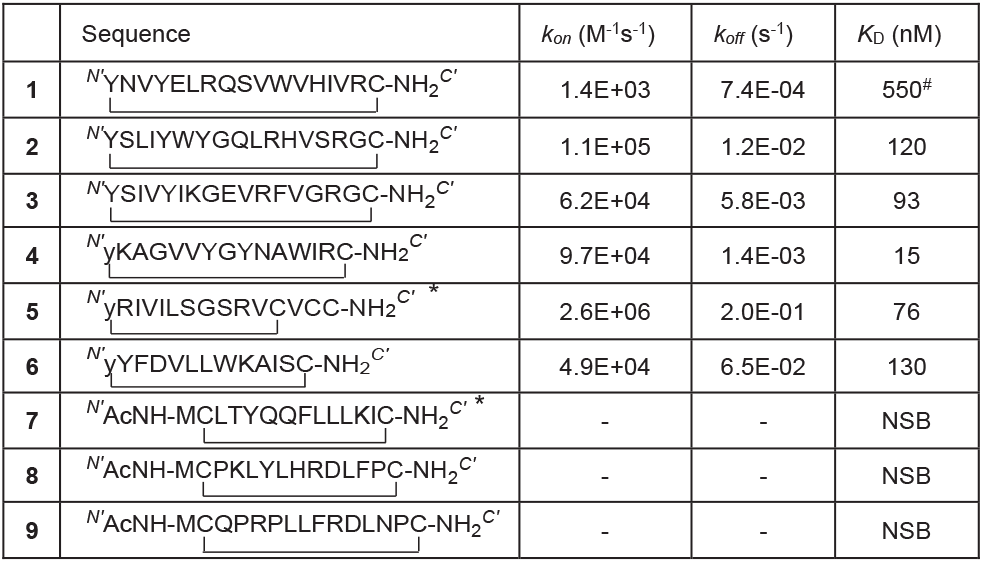
Binding kinetics and dissociation constants (*K*_D_) for the 9 SARS-CoV-2 RBD cyclic peptide ligands synthesized and evaluated against the RBD by SPR. * peptides contain C-terminal hexaethylene glycol moieties; # refractive index values high on sensorgram; NSB = non-specific binding (see Supporting Information for sensorgrams).

The antiviral activity of **4** and **5** was assessed in VeroE6 cells using high content fluorescence microscopy. Briefly, the cyclic peptides and SARS-CoV-2 were pre-incubated for 1 h and then co-incubated with VeroE6 cells for 72 h (MOI 0.5) within 384-well plates. SARS-CoV-2 infection under these conditions leads to viral cytopathic effects (CPE) and cell loss that is correlated to the level of available infectious virus in the initial inoculum. Addition of inhibitors that bind the RBD of the spike protein, thus preventing interactions with cell surface ACE2, e.g. monoclonal antibodies, reduce CPE/cell loss in a dose dependent manner.^20–22^ To enumerate the reduction of CPE in high content, live cell nuclei were stained using Hoechst 33342 (NucBlue) and then the entire 384-well plate imaged (see Supporting Information). Nuclei were then counted in high content using InCarta image analysis software used to give a quantitative measure of cytopathic effect (see Supporting Information). Unfortunately, despite their nM affinities for RBD, neither **4** nor **5** showed antiviral activity against SARS-CoV-2 up to a concentration of 10 μM. Additionally, no inhibition was seen in a SARS-CoV-2 pseudovirion neutralization assay at peptide concentrations of up to 50 μM (see Supporting Information). These data were corroborated by biolayer interferometry (BLI) experiments, in which **4** and **5** were unable to disrupt the interaction between ACE2 and SARS-CoV-2 RBD (see Supporting Information).

Although disappointed with the lack of antiviral activity of the RBD-binding cyclic peptides, we rationalized that the affinity of the peptides could instead prove useful for detection of the spike protein. In order to assess this possibility, we developed an ELISA-based detection assay for the full spike ectodomain and the spike RBD. Peptides **4** and **5** were first re-synthesized bearing a C-terminal biotin tag (see Supporting Information). ELISA plates were first coated with BD-218, a neutralizing monoclonal antibody identified from a COVID-19 patient.^28^ Either spike ectodomain trimer or RBD alone was captured and then detected with biotinylated **4** or **5** and HRP-conjugated streptavidin (Figure 3A). In a second strategy, ELISA plates were directly coated with spike or RBD protein and antigen detected as above (Figure 3B). Gratifyingly, detection of both the RBD and spike ectodomain was possible with biotinylated **4** and **5**, however **5** had substantial non-specific binding, as evidenced by detection of unrelated protein, bovine serum albumin (BSA). In further quantitative assays, biotinylated **4** allowed detection of spike ectodomain trimer at concentrations as low as 30 ng mL^−1^ (see Supporting Information).

**Figure 3.**
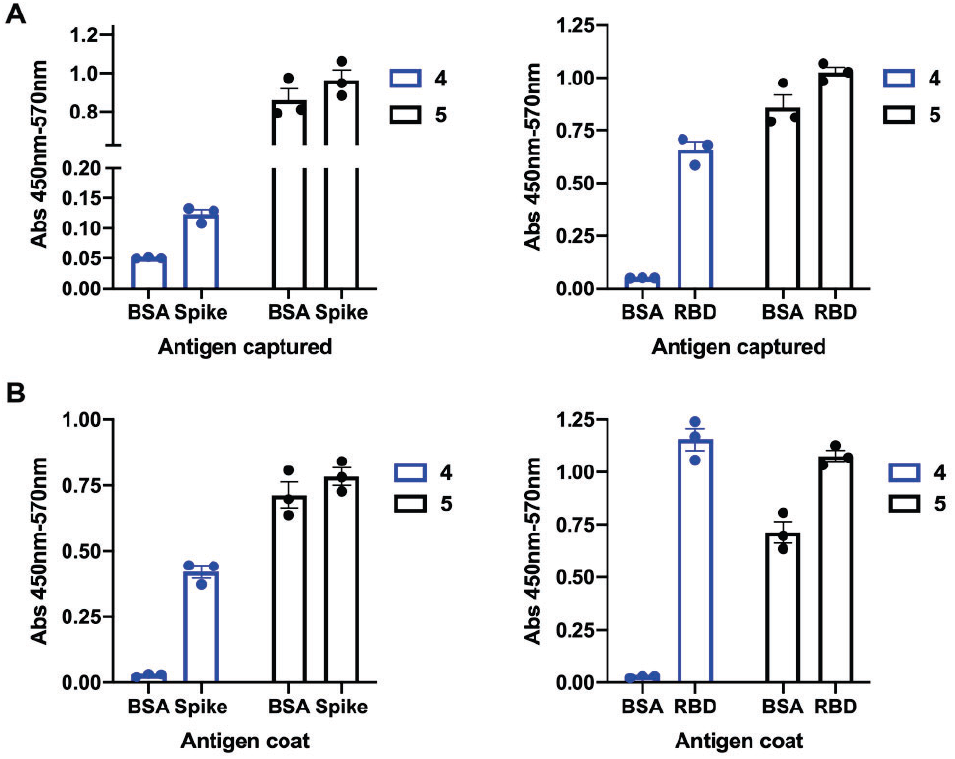
Detection of the spike ectodomain or the spike RBD from SARS-CoV-2 using biotinylated variants of **4** and **5** by ELISA. **A)** Antigen capture ELISA. Plate was coated with antibody BD-218, blocked with BSA, then spike or RBD (1 μg mL^−1^) was captured and detected with biotinylated **4** and **5. B)** Direct ELISA. Plate was coated with spike ectodomain or RBD (1 μg mL^−1^), blocked with BSA, and viral antigen detected with biotinylated **4** and **5**. Individual data points for technical triplicates are shown, with SEM.

Finally, we sought to understand the mechanism by which peptide **4** binds to the RBD. To this end, we obtained crystals of the RBD-**4** complex and solved the structure of the complex by X-ray crystallography to a resolution of 3.96 Å (Figure 4; PDB code 7L4Z). Despite the relatively low resolution, we could identify the binding site of peptide **4,** which lay near the clearly N- and C-termini of the RBD (Figure 4B and Figure S7). Unexpectedly, the peptide binds to a cryptic region of the RBD by displacing a C-terminal β-strand of the domain (^523^TVCG^526^) with residues ^5^VVYG^8^ of the peptide, which adopt essentially the same backbone conformation and make corresponding interactions with the RBD. Interestingly, a similar motif is observed in a linear RBD peptide ligand (^1^TVFG^4^; Figure 4C) that was previously identified by affinity selection-mass spectrometry by Pentelute and co-workers.^44^ Ala mutagenesis of this 13-residue linear peptide highlighted the importance of the motif, particularly residues Val2 and Gly4, which are also present in the sequences of both the RBD C-terminus and peptide **4**. It is therefore tempting to speculate that a similar mode of binding for RBD would be observed for the linear peptide that was discovered through an independent affinity selection method.

**Figure 4.**
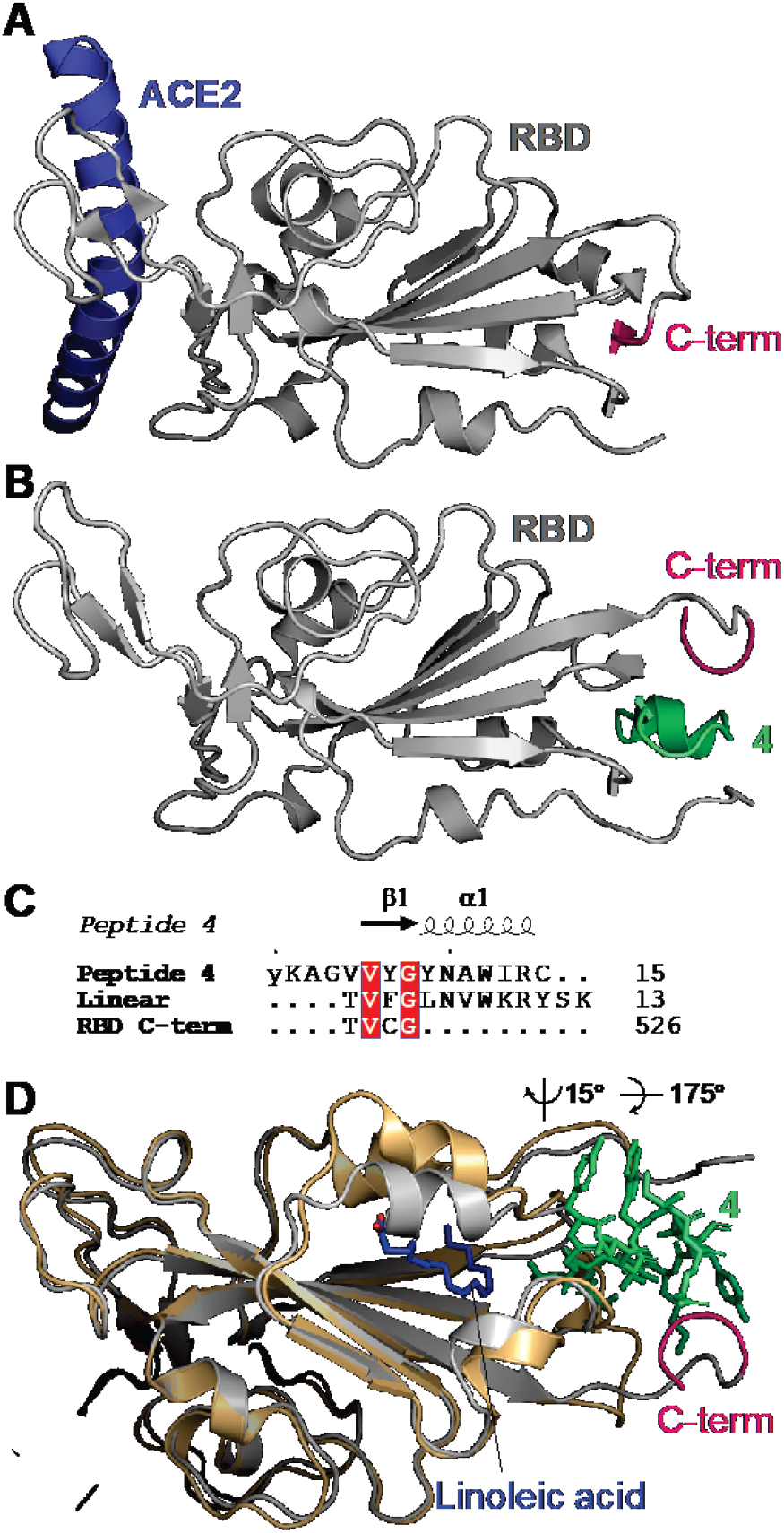
**A)** Structure of ACE2 (only helix 1 shown; blue) in complex with spike RBD (grey; PDB ID 6M0J). The C-terminal β-strand (^523^TVCG^526^) is highlighted in pink. **B)** Structure of spike RBD in complex with peptide **4** (green). **C)** Alignment of Peptide **4**, linear RBD-binding peptide ligand44 and spike RBD C-terminal β-strand. **D)** Structure of RBD-**4** (grey, with **4** shown in green sticks) superimposed onto the spike-linoleic acid complex (PDB code 6ZB4; shown in gold cartoon and blue sticks, respectively). Only the RBD is shown for clarity.

The peptide-binding site is distal from the ACE2-binding interface (Figure 4A), providing a rationale for the lack of antiviral activity. Within the context of the full-length spike protein, peptide binding would only be achieved if the C-terminal β-strand of the RBD undergoes a similar rearrangement to what we observe in our RBD-**4** crystal structure. Moreover, conformational changes in the regions directly flanking the RBD domain would also be required, as the helix formed by peptide **4** residues ^9^YNAWIR^14^ would otherwise clash with these regions. Although peptide **4** could differentially detect spike when the latter was immobilized either by direct adsorption or indirectly via BD-218 antibody capture (Figure 3), we were unable to detect an interaction between **4** and spike using SPR under the conditions tested (see Supporting Information). Together, these data suggest that spike may undergo conformational changes that expose the peptide-binding site when directly immobilized onto plates, and that **4** may therefore be useful in detecting full-length spike using ELISA-based approaches.

## CONCLUSIONS

In summary, we have shown the utility of cyclic peptide mRNA display for the discovery of high affinity ligands to a viral glycoprotein, specifically the spike protein of SARS-CoV-2. We identified six peptides that bind to the RBD of the SARS-CoV-2 spike protein with nanomolar dissociation constants, including two with *K*_D_ values <100 nM. While these peptides were unable to neutralize SARS-CoV-2 infection of cells, we identified a 15residue cyclic peptide (**4**) which was capable of selectively and quantitatively detecting SARS-CoV-2 spike or RBD protein in an ELISA-based detection assay. Moreover, we solved the structure of **4** in complex with SARS-CoV-2 spike RBD. The unexpected binding mode of the cyclic peptide adjacent to the N- and C-termini of the domain suggests that significant structural reorganization of the spike protein would be necessary to accommodate the ligand. The binding of the ligand distal to the ACE2 binding site of the RBD explains the lack of antiviral activity of the cyclic peptide. Importantly, while there have been more than 120 structures of RBD-ligand complexes deposited in the protein data bank over the past 12 months, this represents the first structure of a peptide complexed with the domain, and the first structure of a ligand bound in this cryptic site. This binding mode is likely made possible by the small size of cyclic peptide **4** that enables it to access the compact site on the RBD. A similar binding mode would be difficult to achieve with ACE2 mimics, antibody and antibody fragments that have been the primary focus of RBD- and spike-binding ligands to date. Future work in our laboratories will capitalize on the unique binding mode discovered here by investigating cyclic peptide **4** (and analogues thereof) as easily accessible and cheap reagents for use in rapid, reproducible and simple ELISA-based quantitation of virally expressed SARS-CoV-2 spike protein in patient or environmental samples. Furthermore, the location of this new binding site adjacent to a recently discovered fatty acid binding pocket on the SARS-CoV-2 RBD raises the interesting possibility of designing lipid-linked variants of **4** (Figure 4D).^57^ This hybridization approach may lead to molecules that possess improved affinity for the spike RBD, as well as antiviral activity.

## METHODS

No unexpected or unusually high safety hazards were encountered during the course of this research. All experimental methods can be found in the Supporting Information.

## ASSOCIATED CONTENT

### Supporting Information

The Supporting Information is available free of charge on the ACS Publications website (PDF).

## ACKNOWLEDGMENTS

We would like to acknowledge funding from the National Health and Medical Research Council (Investigator Grant APP1174941 to R.J.P.) and a seed grant from the Drug Discovery Initiative and the Marie Bashir Institute at the University of Sydney. We would also like to thank Dr Luke Dowman for producing Figure 1 included in this manuscript. We thank the beamline scientists at the Australian Synchrotron for their assistance with data collection. This research was undertaken using the MX2 beamline at the Australian Synchrotron, part of ANSTO, and made use of the Australian Cancer Research Foundation (ACRF) detector. Cyclic peptide display screening and SPR was supported by Sydney Analytical, University of Sydney, and sequencing was conducted at the Ramaciotti Centre for Genomics, University of New South Wales.

## ABBREVIATIONS

ACE2: angiotensin converting enzyme 2
BLI: biolayer interferometry
BSA: bovin serum albumin
COVID-19: Coronavirus disease 19
CPE: cytopathic effect(s)
EDC: 1-ethyl-3-(3-dimethylaminopropyl)carbodiimide
ELISA: enzyme-linked immunosorbent assay
*K*_D_: dissociation constant
mRNA: messenger ribonucleic acid
NHS: N-hydroxysuccinimide
PDB: protein data bank
PCR: polymerase chain reaction
RBD: receptor binding domain
SARS-CoV-2: Severe acute respiratory syndrome coronavirus-2
SPR: surface plasmon resonance
TFA: trifluoroacetic acid

## Notes

### Competing Interest Statement

The authors have declared no competing interest.

