## Supporting Information for "Discovery of Cyclic Peptide Ligands to the SARS-CoV-2 Spike Protein using mRNA Display"

### General Procedures

#### General Materials and Methods

Peptide grade *N,N*-dimethylformamide (DMF) for peptide synthesis was purchased from RCI. Gradient grade acetonitrile (MeCN) for chromatography was purchased from Sigma Aldrich and ultrapure water (Type 1) was obtained from a Merck Millipore Direct-Q 5 water purification system. Standard Fmoc-protected amino acids (Fmoc-Xaa-OH), coupling reagents and resins were purchased from Mimotopes or Novabiochem. PEG reagents were obtained from Broadpharm. Fmoc-SPPS was performed manually with these reagents and solvents in polypropylene Teflon-fritted syringes purchased from Torviq or through automated synthesis on a Syro I peptide synthesizer (Biotage). All other reagents were purchased from Sigma Aldrich, AK Scientific or Merck and used as received.

**mRNA-linked cyclic peptide library synthesis.** DNA oligos (Table 1) comprising a T7 promoter, ribosome binding site, ATG start codon, 4–15 NNS (N = A, C, G or T; S = C or G) codons, a TGC (Cys) codon and a 3’ fixed region encoding a Gly-Asn-Leu-Ile linker were amplified by 7 cycles of PCR. The resulting library was transcribed *in vitro* using T7 RNA polymerase to generate mRNA, which was purified by denaturing urea polyacrylamide gel electrophoresis. The libraries of different lengths were then pooled proportional to theoretical diversity as previously described.[1] This final pooled mRNA library was then ligated to a puromycin linked oligonucleotide using T4 RNA ligase. Ribosomal synthesis of the macrocyclic peptide library from the puromycin-linked mRNA library was performed using the PURExpress ΔRF kit (New England Biolabs). With RF2 and RF3 added as per the manufacturer’s instructions. For genetic code reprogramming of the initiating Met residue, a custom “Solution A” was used supplemented with 19 amino acids (-Met) and *N*-chloroacetyl-tyrosine aminoacylated initiator tRNA as previously described.[2] Following translation, the ribosome was denatured by addition of EDTA, and the resulting mRNA-peptide library was reverse transcribed using RNase H- reverse transcriptase.

**Table 1 – Oligonucleotides for library synthesis.**

| | |
|---|---|
| Forward primer | TAATACGACTCACTATAGGGTTAACTTTAAGAAGGAGATATACATA |
| Reverse primer | TTTCCGCCCCCCGTCCTAGATTAAGTTACCGCA |
| MXCGNLI-NNS4.F53 | TTAAGAAGGAGATATACATATGNNSNNSNNSNNSTGCGGTAACTTAATCTAGG |

| | |
|---|---|
| MXCGNLI-NNS5.F56 | TTAAGAAGGAGATATACATATGNNSNNSNNSNNSNNSTGCGGTAACTTAATCTAGG |
| MXCGNLI-NNS6.F59 | TTAAGAAGGAGATATACATATGNNSNNSNNSNNSNNSNNSTGCGGTAACTTAATCTAGG |
| MXCGNLI-NNS7.F62 | TTAAGAAGGAGATATACATATGNNSNNSNNSNNSNNSNNSNNSTGCGGTAACTTAATCTAGG |
| MXCGNLI-NNS8.F65 | TTAAGAAGGAGATATACATATGNNSNNSNNSNNSNNSNNSNNSNNSTGCGGTAACTTAATCTAGG |
| MXCGNLI-NNS9.F68 | TTAAGAAGGAGATATACATATGNNSNNSNNSNNSNNSNNSNNSNNSNNSTGCGGTAACTTAATCTAGG |
| MXCGNLI-NNS10.F71 | TTAAGAAGGAGATATACATATGNNSNNSNNSNNSNNSNNSNNSNNSNNSNNSTGCGGTAACTTAATCTAGG |
| MXCGNLI-NNS11.F74 | TTAAGAAGGAGATATACATATGNNSNNSNNSNNSNNSNNSNNSNNSNNSNNSNNSTGCGGTAACTTAATCTAGG |
| MXCGNLI-NNS12.F77 | TTAAGAAGGAGATATACATATGNNSNNSNNSNNSNNSNNSNNSNNSNNSNNSNNSNNSTGCGGTAACTTAATCTAG<br>G |
| MXCGNLI-NNS13.F80 | TTAAGAAGGAGATATACATATGNNSNNSNNSNNSNNSNNSNNSNNSNNSNNSNNSNNSNNSTGCGGTAACTTAATC<br>TAGG |
| MXCGNLI-NNS14.F83 | TTAAGAAGGAGATATACATATGNNSNNSNNSNNSNNSNNSNNSNNSNNSNNSNNSNNSNNSNNSTGCGGTAACTTA<br>ATCTAGG |
| MXCGNLI-NNS15.F86 | TTAAGAAGGAGATATACATATGNNSNNSNNSNNSNNSNNSNNSNNSNNSNNSNNSNNSNNSNNSNNSTGCGGTAAC<br>TTAATCTAGG |
| MCXCGNLI-NNS4.F56 | TTAAGAAGGAGATATACATATGTGCNNSNNSNNSNNSTGCGGTAACTTAATCTAGG |
| MCXCGNLI-NNS5.F59 | TTAAGAAGGAGATATACATATGTGCNNSNNSNNSNNSNNSTGCGGTAACTTAATCTAGG |
| MCXCGNLI-NNS6.F62 | TTAAGAAGGAGATATACATATGTGCNNSNNSNNSNNSNNSNNSTGCGGTAACTTAATCTAGG |
| MCXCGNLI-NNS7.F65 | TTAAGAAGGAGATATACATATGTGCNNSNNSNNSNNSNNSNNSNNSTGCGGTAACTTAATCTAGG |
| MCXCGNLI-NNS8.F68 | TTAAGAAGGAGATATACATATGTGCNNSNNSNNSNNSNNSNNSNNSNNSTGCGGTAACTTAATCTAGG |
| MCXCGNLI-NNS9.F71 | TTAAGAAGGAGATATACATATGTGCNNSNNSNNSNNSNNSNNSNNSNNSNNSTGCGGTAACTTAATCTAGG |
| MCXCGNLI-NNS10.F74 | TTAAGAAGGAGATATACATATGTGCNNSNNSNNSNNSNNSNNSNNSNNSNNSNNSTGCGGTAACTTAATCTAGG |
| MCXCGNLI-NNS11.F77 | TTAAGAAGGAGATATACATATGTGCNNSNNSNNSNNSNNSNNSNNSNNSNNSNNSNNSTGCGGTAACTTAATCTAG<br>G |
| MCXCGNLI-NNS12.F80 | TTAAGAAGGAGATATACATATGTGCNNSNNSNNSNNSNNSNNSNNSNNSNNSNNSNNSNNSTGCGGTAACTTAATC<br>TAGG |
| MCXCGNLI-NNS13.F83 | TTAAGAAGGAGATATACATATGTGCNNSNNSNNSNNSNNSNNSNNSNNSNNSNNSNNSNNSNNSTGCGGTAACTTA<br>ATCTAGG |
| MCXCGNLI-NNS14.F86 | TTAAGAAGGAGATATACATATGTGCNNSNNSNNSNNSNNSNNSNNSNNSNNSNNSNNSNNSNNSNNSTGCGGTAAC<br>TTAATCTAGG |
| MCXCGNLI-NNS15.F89 | TTAAGAAGGAGATATACATATGTGCNNSNNSNNSNNSNNSNNSNNSNNSNNSNNSNNSNNSNNSNNSNNSTGCGGT<br>AACTTAATCTAGG |

**Affinity selection for SARS-CoV-2 spike RBD.** The reverse transcribed peptide-mRNA library was panned against 200 nM biotinylated SARS-COV-2 RBD immobilized on Dynabeads M-280 streptavidin (Life Technologies) for 30 min at 4 °C. After washing, the fused peptide–mRNA/cDNA was isolated from the beads by heating to 95 °C for 5 min, and cDNA was amplified by PCR, purified by ethanol precipitation and transcribed as above to produce the enriched mRNA library for the next round of selection. For the second and subsequent rounds of selection, three iterative counter-selections were used to remove peptides with affinity for the streptavidin beads. Sequencing of the final enriched cDNA was conducted using an iSeq next generation sequencer (Illumina).

#### Fmoc-Solid-Phase Peptide Synthesis (SPPS)

**General procedure A; Automated Peptide Synthesis (SYRO I peptide synthesizer):** The resin (90 mg, 50 μmol, 0.56 mmol $g^{-1}$, 1 eq.) was treated with 40 vol.% piperidine in DMF (800 μL) for 4 min, drained, then treated with 20 vol.% piperidine in DMF (800 μL) for 4 min, drained, and washed with DMF (4 × 1.2 mL). The resin was then treated with a solution of Fmoc-Xaa-OH or chloroacetic acid (200 μmol, 4 eq.) and Oxyma (220 μmol, 4.4 eq.) in DMF (400 μL), a 1 wt.% solution of 1,3-diisopropyl-2-thiourea in DMF (400 μL), followed by a solution of DIC (200 μmol, 4 eq.) in DMF (400 μL). Coupling of Fmoc-Cys(Trt)-OH and Fmoc-His(Trt)-OH were carried out at 50 ºC for 30 min. All other coupling reactions were conducted at 75 ºC for 15 min. The resin was then drained and washed with DMF (4 × 1.2 mL) before being treated with a solution of 5 vol.% $Ac_2O$ and 10 vol.% *i*$Pr_2NEt$ in DMF (800 μL) for 6 min at room temperature, drained and washed with DMF (4 × 1.2 mL).

For peptides **7–9**, the linear peptide was *N*-terminally acetylated under the following conditions. The resin was treated with 40 vol.% piperidine in DMF (800 μL) for 4 min, drained, then treated with 20 vol.% piperidine in DMF (800 μL) for 4 min, drained, and washed with DMF (4 × 1.6 mL). The resin was then treated with a solution of 5 vol.% $Ac_2O$ and 10 vol.% *i*$Pr_2NEt$ in DMF (800 μL) for 6 min at rt, drained and washed with DMF (4 × 1.2 mL).

**General procedure B; Manual Peptide Synthesis:** The resin (90 mg, 50 μmol, 0.56 mmol $g^{-1}$, 1 eq.) was treated with 20 vol.% piperidine in DMF (5 mL) for 2 × 5 min, filtered, then washed with DMF (5 × 5 mL), $CH_2Cl_2$ (5 × 5 mL) and DMF (5 × 5 mL). The resin was then treated with a solution of Fmoc-Xaa-OH or chloroacetic acid (4 eq.), Oxyma (4.4 eq.) and DIC (4 eq.) in DMF (0.1 M with respect to resin loading) and shaken at 50 ºC for 1 h. The resin was then drained and washed with DMF (5 × 5 mL), $CH_2Cl_2$ (5 × 5 mL) and DMF (5 × 5 mL). The resin was then treated with a solution of 5 vol.% $Ac_2O$ and 10 vol.% *i*$Pr_2NEt$ (10 vol.%) in DMF (2.5 mL) for 3 min at room temperature before being washed with DMF (5 × 5 mL), $CH_2Cl_2$ (5 × 5 mL) and DMF (5 × 5 mL).

**General procedure C: Manual cleavage:** The resin was thoroughly washed with $CH_2Cl_2$ (5 × 5 mL) before being treated with 90:5:5 v/v/v TFA:tri*iso*propylsilane:$H_2O$ and shaken at room temperature for 2 h. The resin was filtered and the filtrate concentrated under a stream of nitrogen before addition of diethyl ether (40 mL). The peptide was pelleted by centrifugation

(4 min, 4 ºC, at 5000 rcf) and the ether was decanted. The crude peptide was dissolved in the minimum volume of 1:1 MeCN/$H_2O$ and concentrated by lyophilization.

**General procedure D: Cyclization 1:** DMSO Cyclization.

The crude peptide (25 µmol) was dissolved in DMSO (5 mL) and *i*$Pr_2NEt$ (180 µL) was added. The peptide solution was heated in a water bath at 60 ºC until the cyclization was complete as judged by UPLC-MS analysis. The solution was then neutralized with the addition of a minimum volume of TFA prior to HPLC purification.

**General procedure E: Cyclization 2:** MeCN/$H_2O$ Cyclization.

The crude peptide (25 µmol) was dissolved in MeCN/$H_2O$ (10 mL) and *i*$Pr_2NEt$ (180 µL) was added. The peptide solution was heated in a water bath at 60 ºC until the cyclization was complete as judged by UPLC-MS analysis. The solution was then neutralized with the addition of a minimum volume of TFA prior to HPLC purification.

**General procedure F: Cyclization 3:** Disulfide Cyclization.

The crude peptide (25 µmol) was dissolved in an aqueous solution of ammonium bicarbonate (50 mL, 50 mM). The mixture was stirred open to the air until the cyclization was complete as judged by UPLC-MS analysis. The solution was concentrated by lyophilization prior to HPLC purification.

**Preparative Liquid Chromatography**

Preparative and semi-preparative reversed-phase high performance liquid chromatography (HPLC) was performed using a Waters 600E multisolvent delivery system with a Rheodyne 7725i injection valve (5 mL loading loop) with a Waters 500 pump and a Waters 490E programmable wavelength detector operating at 214 nm and 280 nm. Preparative reversed-phase HPLC was performed using a Waters X-Bridge® C18 OBD™ Prep Column (5 µm, 30 × 150 mm) at a flow rate of 38 mL $min^{-1}$ using a mobile phase of 0.1% TFA in water (solvent A) and 0.1% TFA in MeCN (solvent B) on linear gradients, unless otherwise specified. Semi-preparative chromatography was performed using a Waters X-Bridge® BEH C18 OBD™ Prep Column (300 Å, 5 µm, 10 × 250 mm) at a flow rate of 4 mL $min^{-1}$ using a mobile phase of 0.1% TFA in water (solvent A) and 0.1% TFA in MeCN (solvent B) on linear gradients, unless otherwise specified.

**Liquid Chromatography-Mass Spectrometry**

Liquid Chromatography-Mass Spectrometry (LC-MS) was performed on a Shimadzu 2020 UPLC-MS instrument with a Nexera X2 LC-30AD pump, Nexera X2 SPD-M30A UV/Vis diode array detector and a Shimadzu 2020 (ESI) mass spectrometer operating in positive ion mode. Separations were performed on a Waters Acquity BEH300 1.7 μm, 2.1 × 50 mm (C18) column at a flow rate of 0.6 mL $min^{-1}$. All separations were performed using a mobile phase of 0.1 vol.% formic acid in water (solvent A) and 0.1 vol.% formic acid in MeCN (solvent B) using linear gradients over 5 min.

Analytical reversed-phase HPLC was performed on a Waters Alliance e2695 HPLC system equipped with a 2998 PDA detector (λ = 210–400 nm). Separations were performed on a Waters XBridge® Peptide BEH300 5 μm, 4.6 × 250 mm (C18) column at 40 ºC with a flow rate of 1.0 mL $min^{-1}$. All separations were performed using a mobile phase of 0.1% TFA in water (Solvent A) and 0.1% TFA in MeCN (Solvent B) using linear gradients, unless otherwise specified.

**Mass Spectrometry**

Low resolution mass spectra were recorded on a Shimadzu 2020 (ESI) mass spectrometer operating in positive and negative mode. High resolution mass spectra were recorded on a Bruker-Daltronics Apex Ultra 7.0 T Fourier transform (FTICR) mass spectrometer.

**Fmoc-amino acids and carboxylic acid derivatives used**

Unless otherwise specified the following carboxylic acids (and derivatives) were used in the assembly of the linear peptides.

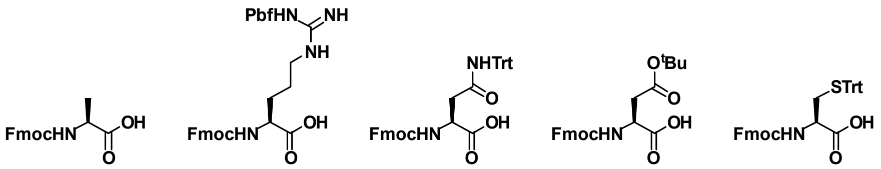

Alanine Ala, A | Arginine Arg, R | Asparagine Asn, N | Aspartic acid Asp, D | Cysteine Cys, C

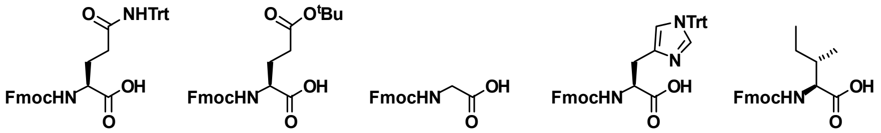

Glutamine Gln, Q | Glutamic acid Glu, E | Glycine Gly, G | Histidine His, H | Isoleucine Ile, I

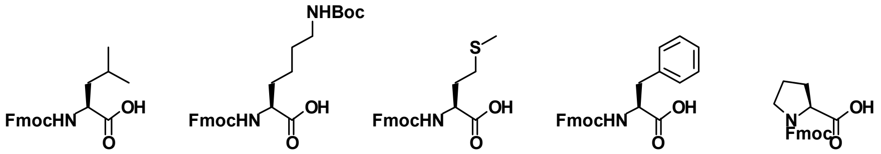

Leucine Leu, L | Lysine Lys, K | Methionine Met, M | Phenylalanine Phe, F | Proline Pro, P

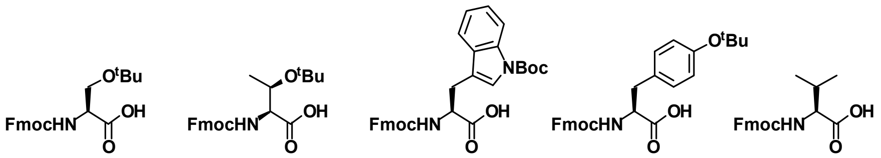

Serine Ser, S | Threonine Thr, T | Tryptophan Trp, W | Tyrosine Tyr, Y | Valine Val, V

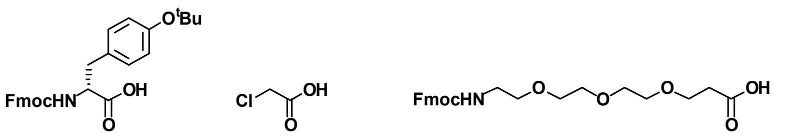

D-Tyrosine D-Tyr, y | Chloroacetic acid | NHFmoc-[PEG$_3$]-OH

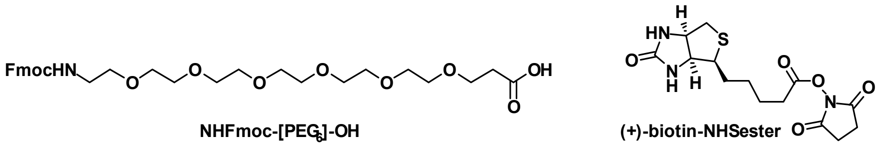

NHFmoc-[PEG$_6$]-OH | (+)-biotin-NHSester

**Synthesis of peptides**

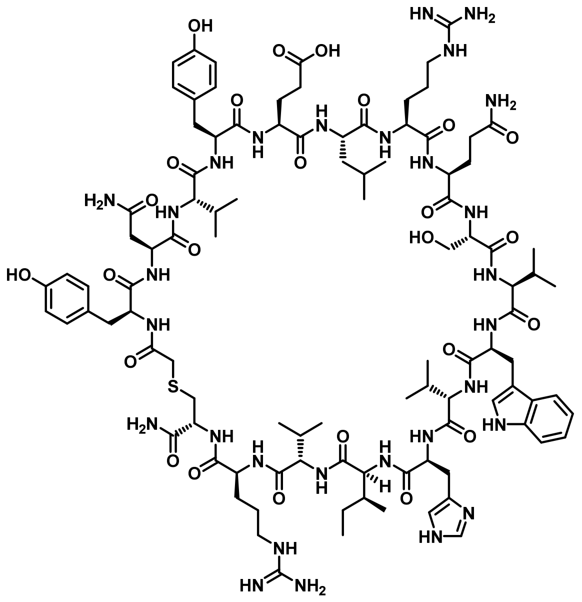

**Peptide 1:** Rink amide resin (90 mg, 50 μmol, 0.56 mmol $g^{-1}$) was loaded with Fmoc-Cys(Trt)-OH according to general procedure **A**. The target peptide was then synthesized by iterative Fmoc solid-phase peptide synthesis (Fmoc SPPS) on the Syro I peptide synthesizer according to general procedure **A**, with chloroacetic acid as the final coupling step to generate the resin-bound sequence YNVYELRQSVWVHIVRC. A portion (25 μmol) of the peptide was cleaved from resin by general procedure **C** and cyclized according to general procedure **D**. Purification by preparative HPLC (5 vol.% MeCN in $H_2O$ with 0.1 vol.% TFA over 10 min then 5 to 40 vol.% MeCN in $H_2O$ with 0.1 vol.% TFA over 50 min), followed by lyophilization, afforded the pure peptide as a white fluffy solid (1.1 mg, 2%). **HRMS:** Calculated for $[C_{101}H_{151}N_{29}O_{25}S+2H]^{2+}$: 1102.06511, found: 1102.06574. **LRMS:** (+ESI) *m/z* 1102.8 $[M+2H]^{2+}$, 735.6 $[M+3H]^{3+}$. **Analytical HPLC:** $R_t$ = 14.9 min (1 to 80 vol.% MeCN in $H_2O$ with 0.1 vol.% TFA over 30 min, λ = 214 nm).

A) **Analytical HPLC Trace** (λ= 214 nm)

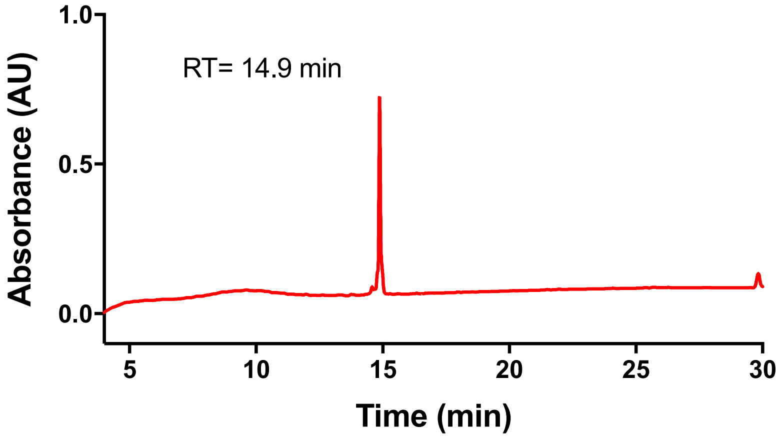


**B) ESI MS (+)**

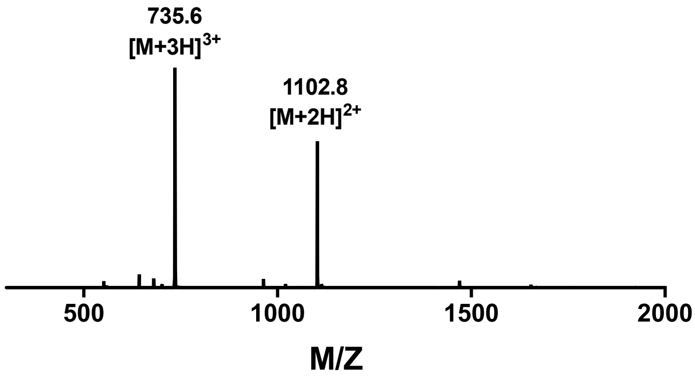

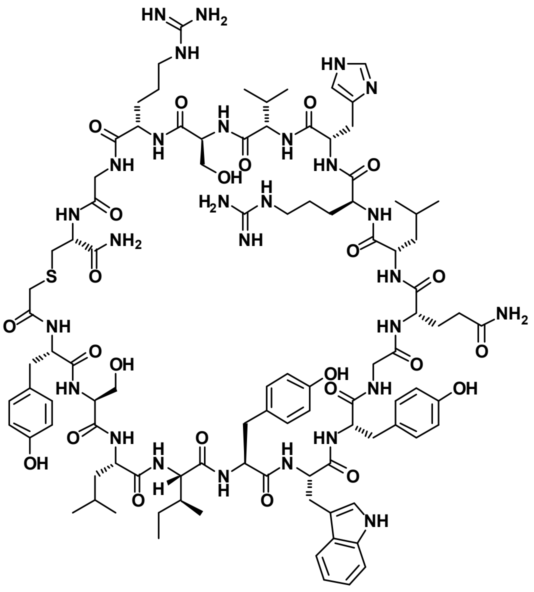


**Peptide 2:** Rink amide resin (90 mg, 50 µmol, 0.56 mmol $g^{-1}$) was loaded with Fmoc-Cys(Trt)-OH according to general procedure **A**. The peptide was then synthesized by iterative Fmoc solid-phase peptide synthesis (Fmoc SPPS) on the Syro I peptide synthesizer according to general procedure **A**, with chloroacetic acid as the final coupling step to generate the resin-bound sequence YSLIYWYGQLRHVSRGC. A portion (25 µmol) of the peptide was cleaved from resin by general procedure **C** and cyclized according to general procedure **D**. Purification by preparative HPLC (5 vol.% MeCN in $H_2O$ with 0.1 vol.% TFA over 10 min then 5 to 55 vol.% MeCN in $H_2O$ with 0.1 vol.% TFA over 50 min), followed by lyophilization, afforded the pure peptide as a white fluffy solid (5.9 mg, 11%). **HRMS:** Calculated for $[C_{99}H_{142}N_{28}O_{24}S+2H]^{2+}$: 1070.53090, found: 1070.53156. **LRMS:** (+ESI) *m/z* 1427.8 $[2M+3H]^{3+}$, 1071.1 $[M+2H]^{2+}$, 714.3 $[M+3H]^{3+}$. **Analytical HPLC:** $R_t$ = 15.5 min (1 to 80 vol.% MeCN in $H_2O$ with 0.1 vol.% TFA over 30 min, λ = 214 nm).

A) **Analytical HPLC Trace** (λ= 214 nm)

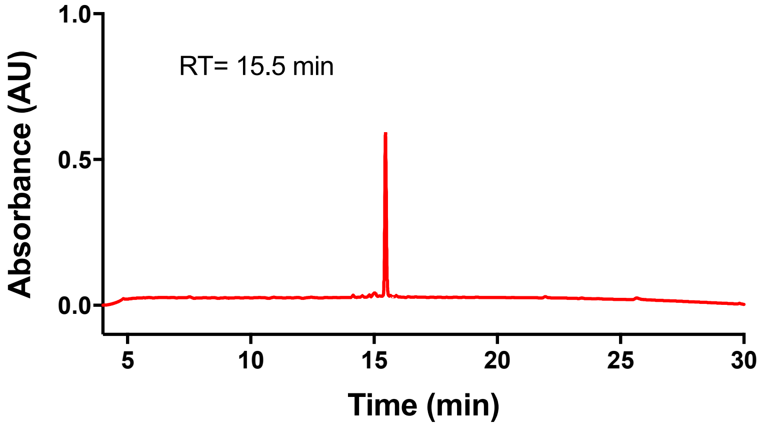


B) **ESI MS (+)**

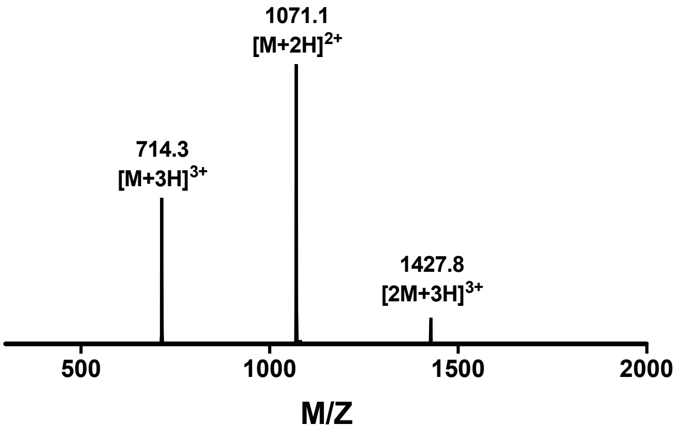

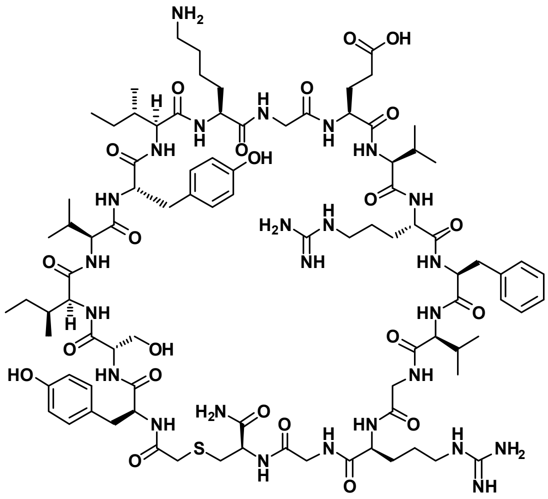


**Peptide 3:** Rink amide resin (90 mg, 50 µmol, 0.56 mmol $g^{-1}$) was loaded with Fmoc-Cys(Trt)-OH according to general procedure **A**. The peptide was then synthesized by iterative Fmoc solid-phase peptide synthesis (Fmoc SPPS) on the Syro I peptide synthesizer according to general procedure **A**, with chloroacetic acid as the final coupling step to generate the resin-bound sequence YSIVYIKGEVRFVGRGC. A portion (25 µmol) of the peptide was cleaved from resin by general procedure **C** and cyclized according to general procedure **D**. Purification by preparative HPLC (5 vol.% MeCN in $H_2O$ with 0.1 vol.% TFA over 10 min then 5 to 50 vol.% MeCN in $H_2O$ with 0.1 vol.% TFA over 50 min), followed by lyophilization, afforded the pure peptide as a white fluffy solid (1.6 mg, 3%). **HRMS:** Calculated for $[C_{91}H_{143}N_{25}O_{23}S+2H]^{2+}$: 993.02492, found: 993.02528. **LRMS:** (+ESI) *m/z* 1986.7 $[M+H]^{+}$, 993.7 $[M+2H]^{2+}$, 662.8 $[M+3H]^{3+}$. **Analytical HPLC:** $R_t$ = 14.4 min (1 to 80 vol.% MeCN in $H_2O$ with 0.1 vol.% TFA over 30 min, λ = 214 nm).

A) Analytical HPLC Trace (λ= 214 nm)

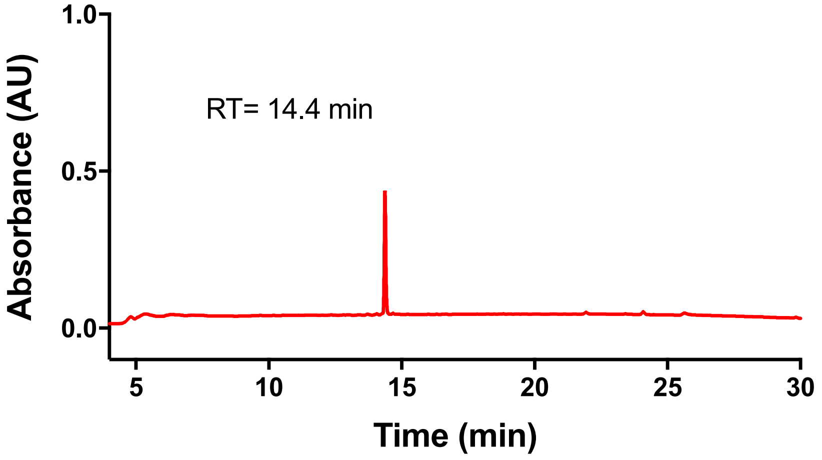


B) ESI MS (+)

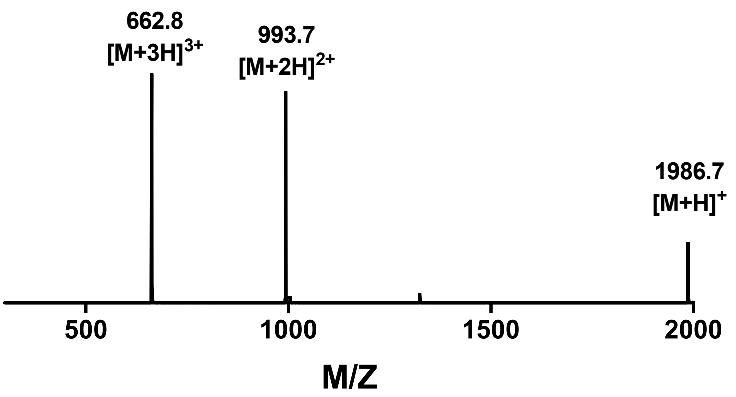

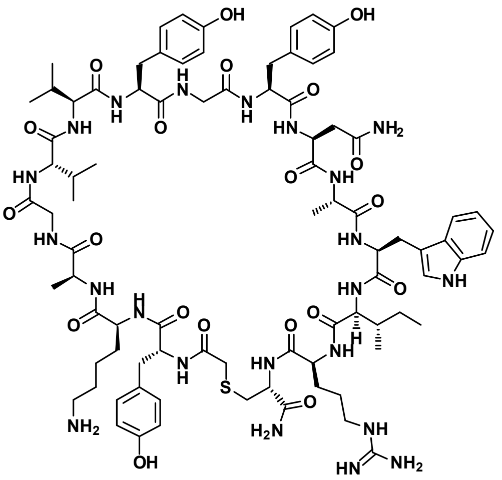

**Peptide 4:** Rink amide resin (90 mg, 50 μmol, 0.56 mmol $g^{-1}$) was loaded with Fmoc-Cys(Trt)-OH according to general procedure **A**. The peptide was then synthesized by iterative Fmoc solid-phase peptide synthesis (Fmoc SPPS) on the Syro I peptide synthesizer according to general procedure **A**, with chloroacetic acid as the final coupling step to generate the resin-bound sequence yKAGVVYGYNAWIRC. A portion (25 μmol) of the peptide was cleaved from resin by general procedure **C** and cyclized according to general procedure **E**. Purification by preparative HPLC (100% $H_2O$ with 0.1 vol.% TFA over 10 min then 0 to 50 vol.% MeCN in $H_2O$ with 0.1 vol.% TFA over 45 min), followed by lyophilization, afforded the pure peptide as a white fluffy solid (2.4 mg, 5%). **HRMS:** Calculated for $[C_{85}H_{120}N_{22}O_{20}S+2H]^{2+}$: 901.44577, found: 901.44457. **LRMS:** (+ESI) *m/z* 1802.7 $[M+H]^{+}$, 901.8 $[M+2H]^{2+}$, 601.5 $[M+3H]^{3+}$. **Analytical HPLC:** $R_t$ = 14.9 min (1 to 80 vol.% MeCN in $H_2O$ with 0.1 vol.% TFA over 30 min, λ = 214 nm).

A) Analytical HPLC Trace (λ= 214 nm)

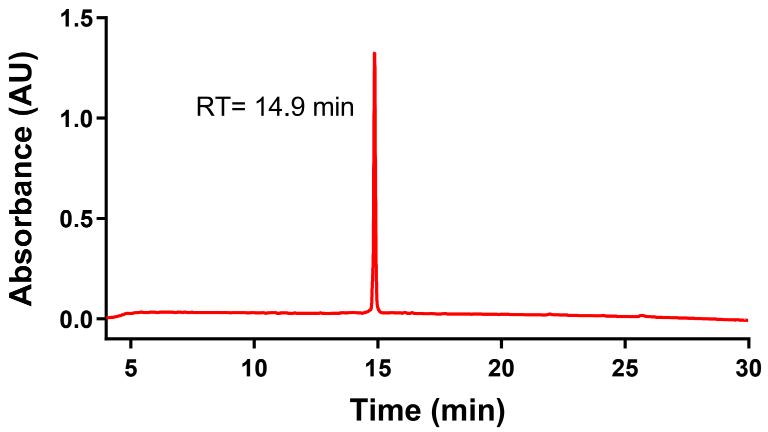


B) ESI MS (+)

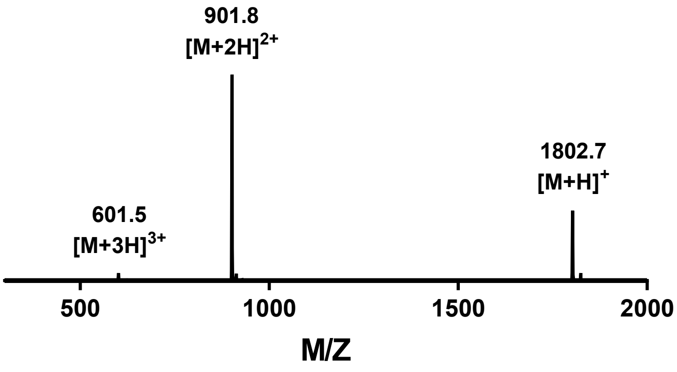

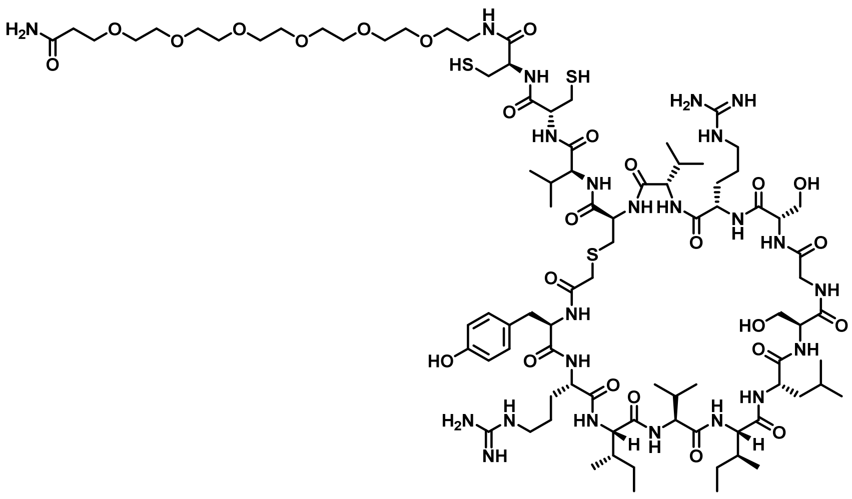

**Peptide 5:** Rink amide resin (90 mg, 50 µmol, 0.56 mmol $g^{-1}$) was loaded with Fmoc-PEG6-OH according to general procedure **A**. The peptide was then synthesized by iterative Fmoc solid-phase peptide synthesis (Fmoc SPPS) on the Syro I peptide synthesizer according to general procedure **A**, with chloroacetic acid as the final coupling step to generate the resin-bound sequence yRIVILSGSRVCVC(Acm)C(Acm)-[$PEG_6$]. Notably, the non-standard Fmoc-protected amino acid, Fmoc-Cys(Acm)-OH, was used as the Fmoc-Xaa-OH for two cysteine residues to enable selective cyclization of the Cys residue closest to the *N*-terminus. A portion (25 µmol) of the peptide was cleaved from resin by general procedure **C** and cyclized according to general procedure **D**. Purification by preparative HPLC (100% $H_2O$ with 0.1 vol.% TFA over 5 min then 0 to 50 vol.% MeCN in $H_2O$ with 0.1 vol.% TFA over 45 min), followed by lyophilization, afforded the Acm-protected peptide as a white fluffy solid. To a solution of the peptide (20 mg, 8.4 µmol) in 1:1 MeCN/$H_2O$ (5 mL) was added silver acetate (700 mg, 4.2 mmol, 500 eq.). The resulting suspension was shaken at room temperature for 1 h, before dithiothreitol (710 mg, 4.6 mmol, 550 eq.) was added. The suspension was centrifuged (5 min, 22 ºC, at 5000 rcf), collecting the supernatant. The pellet was reagitated with DMSO (5 mL) and centrifuged (5 min, 22 ºC, at 5000 rcf), collecting the supernatant. The combined supernatants were purified by preparative HPLC (100% $H_2O$ with 0.1 vol.% TFA over 10 min then 0 to 50 vol.% MeCN in $H_2O$ with 0.1 vol.% TFA over 45 min), followed by lyophilization, afforded the pure peptide as a white fluffy solid (2.0 mg, 4%). **HRMS:** Calculated for $[C_{88}H_{153}N_{23}O_{26}S_3+2H]^{2+}$: 1022.02541, found: 1022.02375. **LRMS:** (+ESI) *m/z* 1364.4 $[2M+3H]^{3+}$, 1023.6 $[M+2H]^{2+}$, 682.7 $[M+3H]^{3+}$. **Analytical HPLC:** $R_t$ = 16.1 min (1 to 80 vol.% MeCN in $H_2O$ with 0.1 vol.% TFA over 30 min, λ = 214 nm).

**A) Analytical HPLC Trace** (λ= 214 nm)

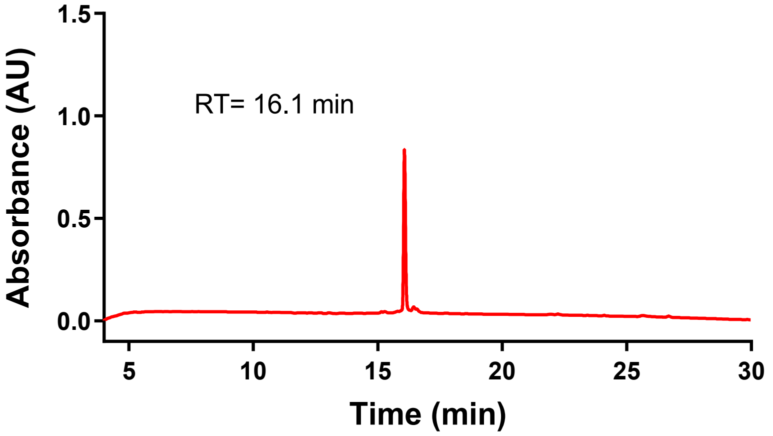


**B) ESI MS (+)**

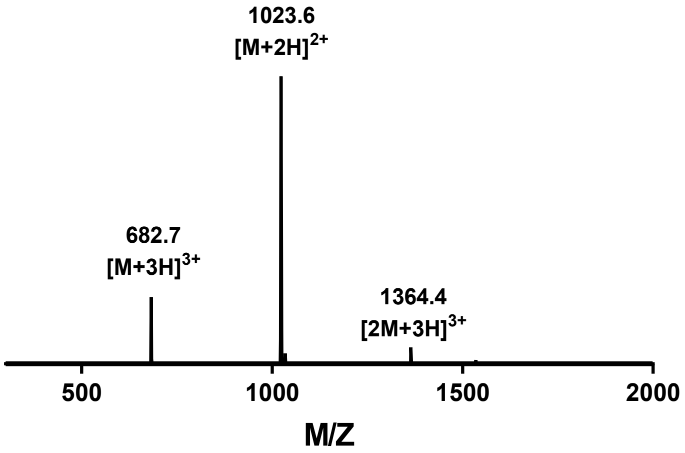

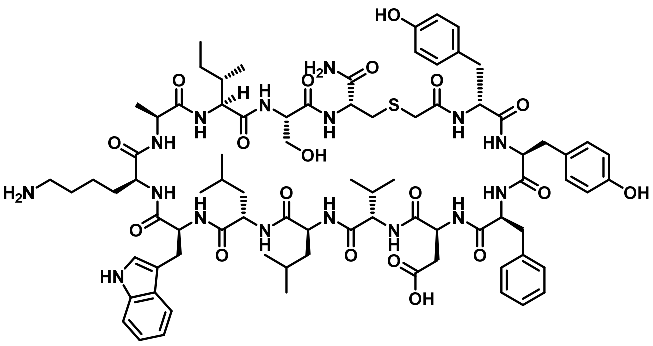


**Peptide 6:** Rink amide resin (90 mg, 50 μmol, 0.56 mmol $g^{-1}$) was loaded with Fmoc-Cys(Trt)-OH according to general procedure **A**. The peptide was then synthesized by iterative Fmoc solid-phase peptide synthesis (Fmoc SPPS) on the Syro I peptide synthesizer according to general procedure **A**, with chloroacetic acid as the final coupling step to generate the resin-bound sequence yYFDVLLWKAISC. A portion (25 μmol) of the peptide was cleaved from resin by general procedure **C** and cyclized according to general procedure **D**. Purification by preparative HPLC (5 vol.% MeCN in $H_2O$ with 0.1 vol.% TFA over 10 min then 5 to 70 vol.% MeCN in $H_2O$ with 0.1 vol.% TFA over 50 min), followed by lyophilization, afforded the pure peptide as a white fluffy solid (5.9 mg, 14%). **HRMS:** Calculated for $[C_{82}H_{116}N_{16}O_{19}S+2H]^{2+}$: 830.41562, found: 830.41587. **LRMS:** (+ESI) *m/z* 1660.6 $[M+H]^+$, 830.9 $[M+2H]^{2+}$. **Analytical HPLC:** $R_t$ = 20.7 min (1 to 80 vol.% MeCN in $H_2O$ with 0.1 vol.% TFA over 30 min, λ = 214 nm).

A) Analytical HPLC Trace (λ= 214 nm)

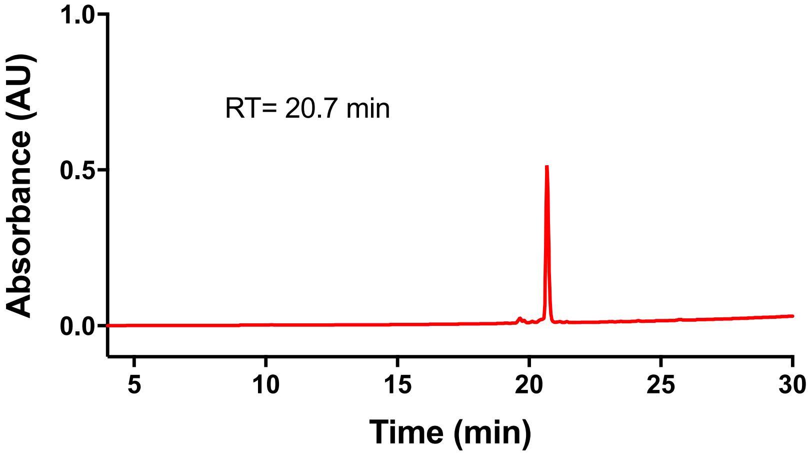


B) ESI MS (+)

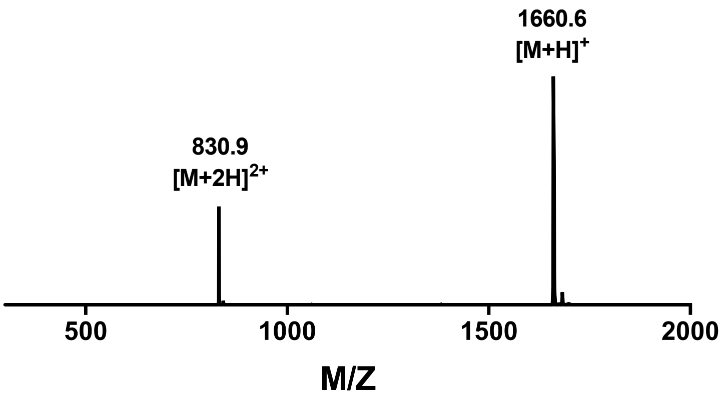

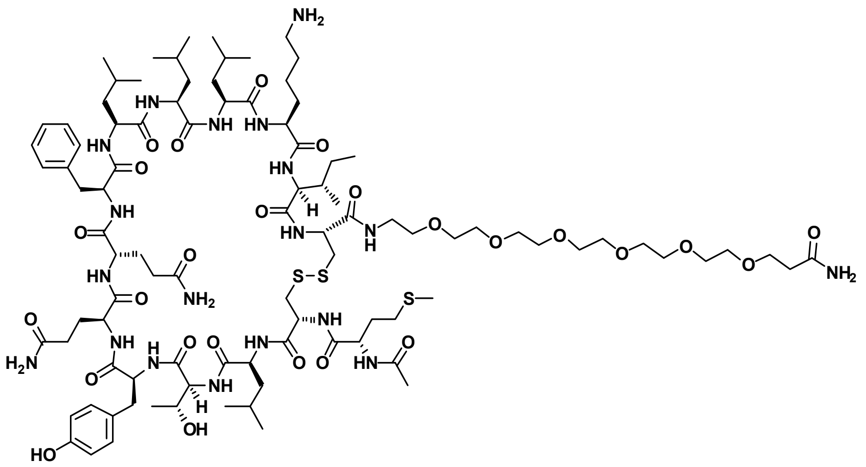

**Peptide 7:** Rink amide resin (90 mg, 50 µmol, 0.56 mmol $g^{-1}$) was loaded with Fmoc-PEG6-OH according to general procedure **B**. The peptide was then synthesized by iterative manual Fmoc solid-phase peptide synthesis (Fmoc SPPS) according to general procedure **B**, with Fmoc-Met-OH as the final coupling step to generate the resin-bound sequence MCLTYQQFLLLKIC. A portion (25 µmol) of the peptide was cleaved from resin by general procedure **C** and oxidatively cyclized according to general procedure **F**. Purification by preparative HPLC (5 vol.% MeCN in $H_2O$ with 0.1 vol.% TFA over 10 min then 5 to 60 vol.% MeCN in $H_2O$ with 0.1 vol.% TFA over 50 min), followed by lyophilization, afforded the pure peptide as a white fluffy solid (1.7 mg, 3%). **HRMS:** Calculated for $[C_{96}H_{161}N_{19}O_{26}S_3+2H]^{2+}$: 1046.05057, found: 1046.05167. **LRMS:** (+ESI) *m/z* 1046.6 $[M+2H]^{2+}$, 698.1 $[M+3H]^{3+}$. **Analytical HPLC:** $R_t$ = 19.8 min (1 to 80 vol.% MeCN in $H_2O$ with 0.1 vol.% TFA over 30 min, $\lambda$ = 214 nm).

A) Analytical HPLC Trace (λ= 214 nm)

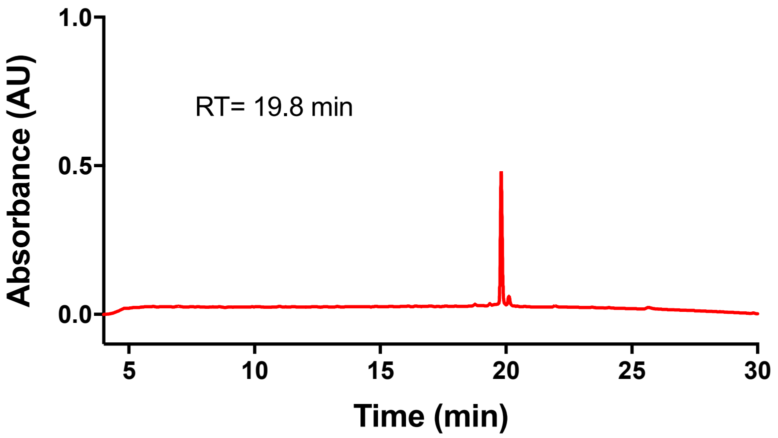


B) ESI MS (+)

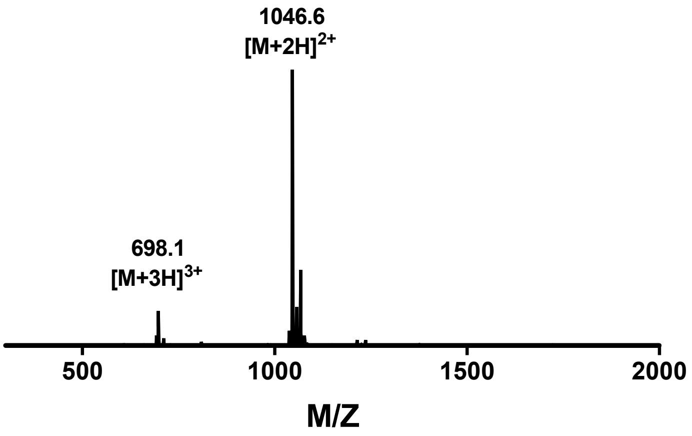

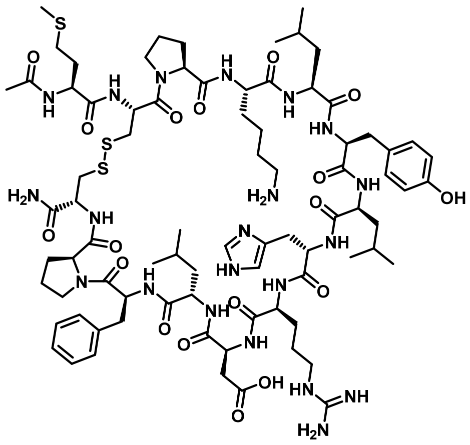


**Peptide 8:** Rink amide resin (90 mg, 50 μmol, 0.56 mmol $g^{-1}$) was loaded with Fmoc-Cys(Trt)-OH according to general procedure **A**. The peptide was then synthesized by iterative Fmoc solid-phase peptide synthesis (Fmoc SPPS) on the Syro I peptide synthesizer according to general procedure **A**, with Fmoc-Met-OH as the final coupling step to generate the resin-bound sequence MCPKLYLHRDLFPC. A portion (25 μmol) of the peptide was cleaved from resin by general procedure **C** and oxidatively cyclized according to general procedure **F**. Purification by preparative HPLC (5 vol.% MeCN in $H_2O$ with 0.1 vol.% TFA over 10 min then 5 to 45 vol.% MeCN in $H_2O$ with 0.1 vol.% TFA over 50 min), followed by lyophilization, afforded the pure peptide as a white fluffy solid (2.6 mg, 6%). **HRMS:** Calculated for $[C_{81}H_{125}N_{21}O_{18}S_3+2H]^{2+}$: 887.93313, found: 887.93379. **LRMS:** (+ESI) *m/z* 1775.9 $[M+H]^+$, 888.4 $[M+2H]^{2+}$, 592.6 $[M+3H]^{3+}$. **Analytical HPLC:** $R_t$ = 16.0 min (1 to 80 vol.% MeCN in $H_2O$ with 0.1 vol.% TFA over 30 min, λ = 214 nm).

A) Analytical HPLC Trace (λ= 214 nm)

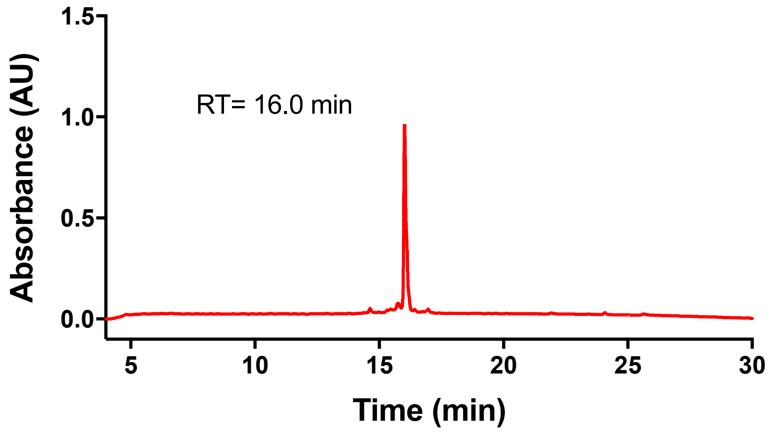


B) ESI MS (+)

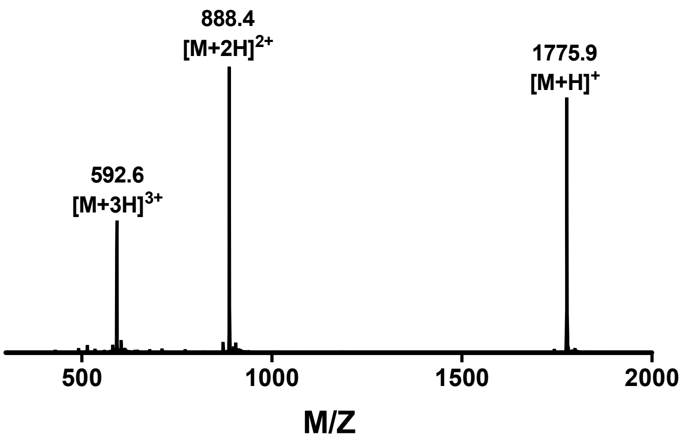

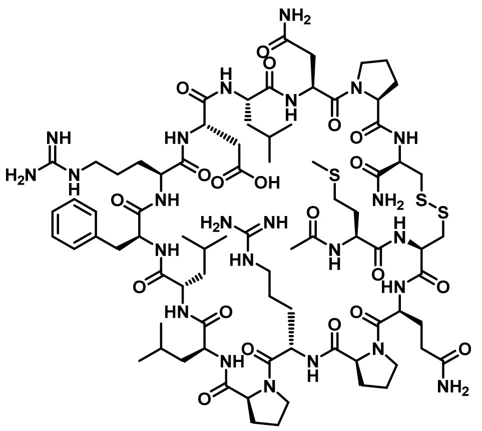

**Peptide 9:** Rink amide resin (90 mg, 50 µmol, 0.56 mmol $g^{-1}$) was loaded with Fmoc-Cys(Trt)-OH according to general procedure **A**. The peptide was then synthesized by iterative Fmoc solid-phase peptide synthesis (Fmoc SPPS) on the Syro I peptide synthesizer according to general procedure **A**, with Fmoc-Met-OH as the final coupling step to generate the resin-bound sequence MCQPRPLLFRDLNPC. A portion (25 µmol) of the peptide was cleaved from resin by general procedure **C** and oxidatively cyclized according to general procedure **F**. Purification by preparative HPLC (5 vol.% MeCN in $H_2O$ with 0.1 vol.% TFA over 10 min then 5 to 45 vol.% MeCN in $H_2O$ with 0.1 vol.% TFA over 50 min), followed by lyophilization, afforded the pure peptide as a white fluffy solid (2.4 mg, 5%). **HRMS:** Calculated for $[C_{80}H_{130}N_{24}O_{20}S_3+2H]^{2+}$: 921.45222, found: 921.45224. **LRMS:** (+ESI) *m/z* 1842.9 $[M+H]^+$, 921.9 $[M+2H]^{2+}$, 614.9 $[M+3H]^{3+}$. **Analytical HPLC:** $R_t$ = 16.3 min (1 to 80 vol.% MeCN in $H_2O$ with 0.1 vol.% TFA over 30 min, λ = 214 nm).

A) Analytical HPLC Trace (λ= 214 nm)

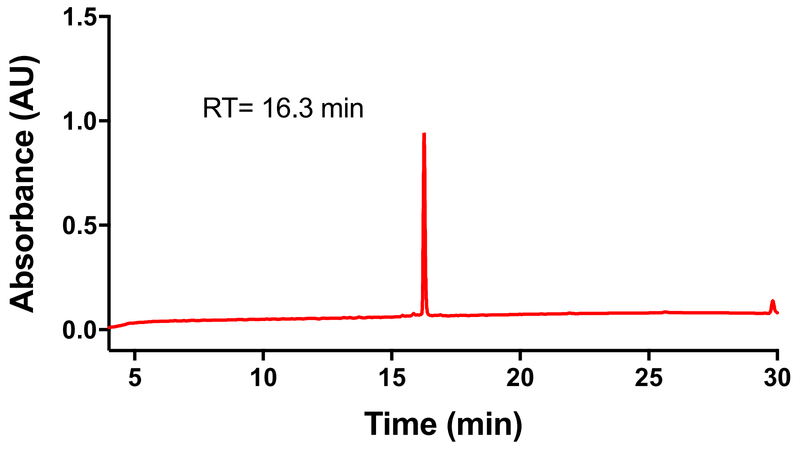


B) ESI MS (+)

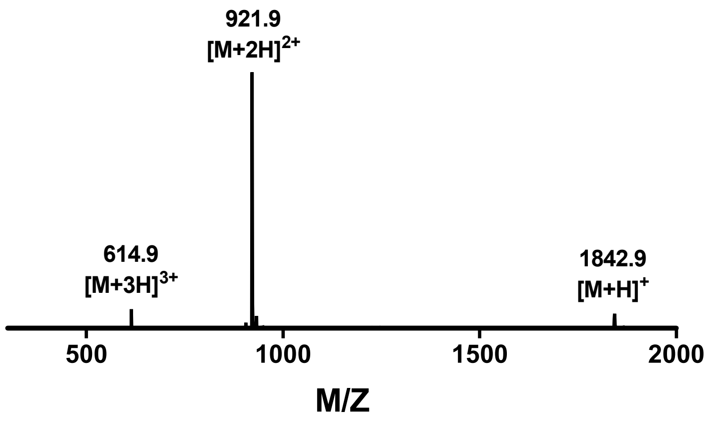

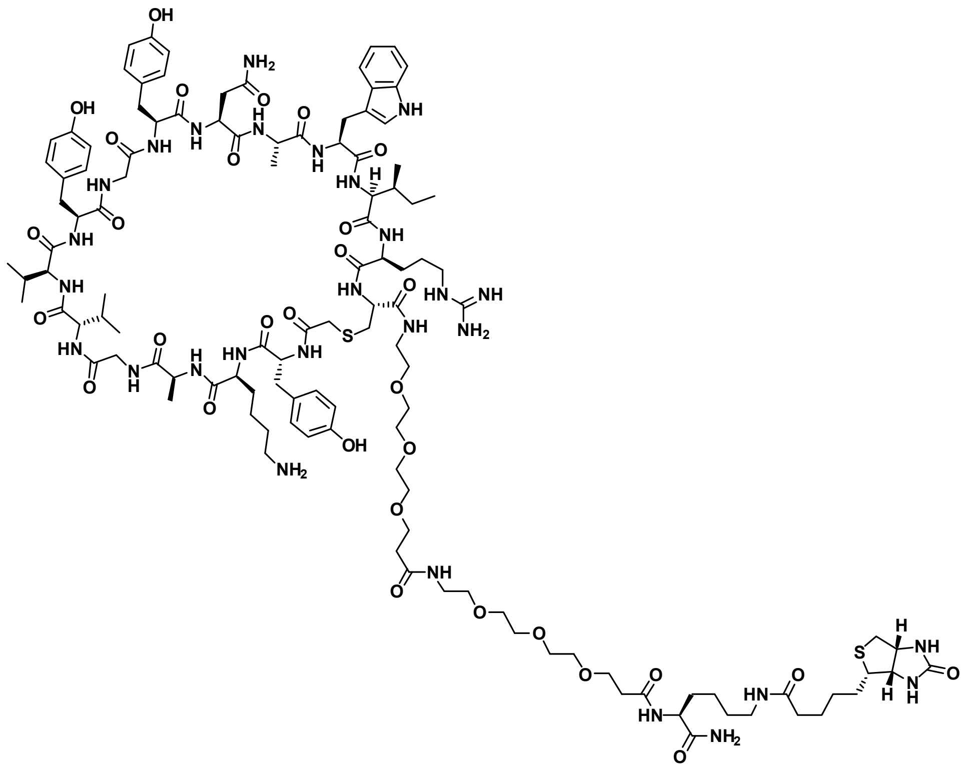

**Biotinylated 4:** Rink amide resin (90 mg, 50 μmol, 0.56 mmol $g^{-1}$) was loaded with Fmoc-Lys(Alloc)-OH according to general procedure **A**. The peptide was then synthesized by iterative Fmoc solid-phase peptide synthesis (Fmoc SPPS) on the Syro I peptide synthesizer according to general procedure **A**, with chloroacetic acid as the final coupling step to generate the resin-bound sequence yKAGVVYGYNAWIRC-[$PEG_3$]-[$PEG_3$]-K(Alloc). A portion (25 μmol) of the peptide was treated with $Pd(PPh_3)_4$ (5 mg, 5 μmol, 0.2 eq.) and $PhSiH_3$ (62 μL, 500, μmol, 20 eq.) in $CH_2Cl_2$ (0.75 mL, 0.03 M with respect to resin loading) and shaken at room temperature for 20 min. The resin was then drained and washed with $CH_2Cl_2$ (10 × 5 mL) and DMF (5 × 5 mL). The resin was then treated with a solution of (+)-biotin *N*-hydroxysuccinimide ester (17 mg, 50 μmol, 2 eq.) and *i*$Pr_2NEt$ (17 μL, 100 μmol, 4 eq.) in DMF (0.25 mL, 0.1 M with respect to resin loading) and shaken at room temperature for 16 h. The resin was then drained and washed with DMF (5 × 5 mL), $CH_2Cl_2$ (5 × 5 mL) and DMF (5 × 5 mL). The resin was then treated with a solution of 5 vol.% $Ac_2O$ and 10 vol.% *i*$Pr_2NEt$ (10 vol.%) in DMF (2.5 mL) for 3 min at room temperature before being washed with DMF (5 × 5 mL), $CH_2Cl_2$ (5 × 5 mL), DMF (5 × 5 mL) and $CH_2Cl_2$ (5 × 5 mL). The peptide was cleaved from resin by general procedure **C** and cyclized according to general procedure **D**. Purification by preparative HPLC (100% $H_2O$ with 0.1 vol.% TFA over 10 min then 0 to 50 vol.% MeCN in $H_2O$ with 0.1 vol.% TFA over 50 min), followed by lyophilization, afforded the pure peptide

as a white fluffy solid (16.5 mg, 24%). **HRMS:** Calculated for $[C_{119}H_{180}N_{28}O_{31}S_2+3H]^{3+}$: 854.76763, found: 854.76792. **LRMS:** (+ESI) *m/z* 1923.3 $[3M+4H]^{4+}$, 1709.6 $[2M+3H]^{3+}$, 1282.4 $[M+2H]^{2+}$, 855.2 $[M+3H]^{3+}$, 641.7 $[M+4H]^{4+}$. **Analytical HPLC:** $R_t$ = 15.1 min (1 to 80 vol.% MeCN in $H_2O$ with 0.1 vol.% TFA over 30 min, λ = 214 nm).

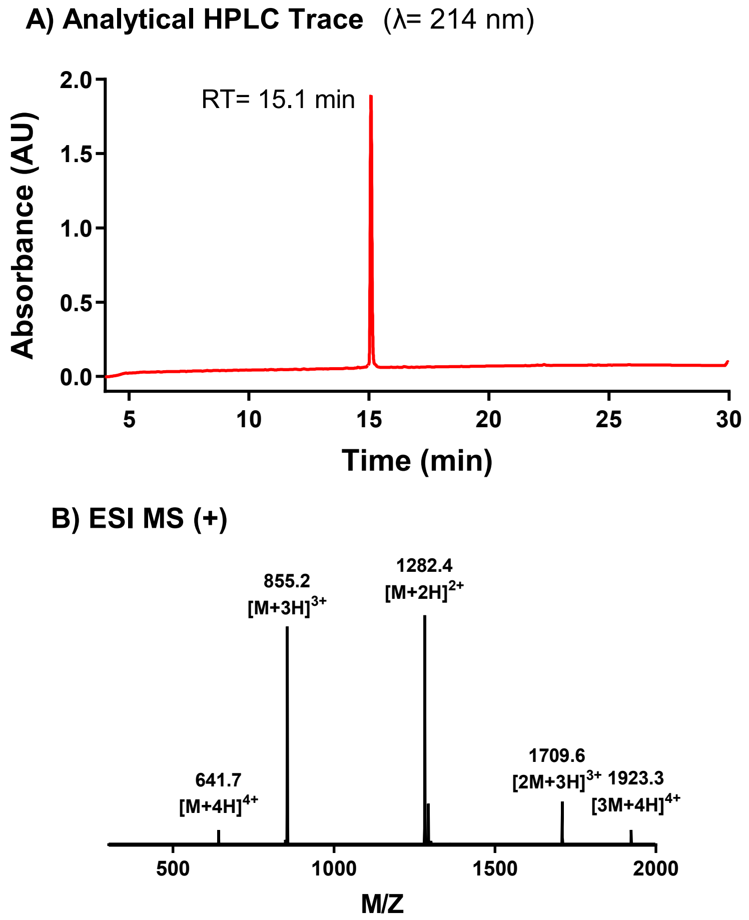

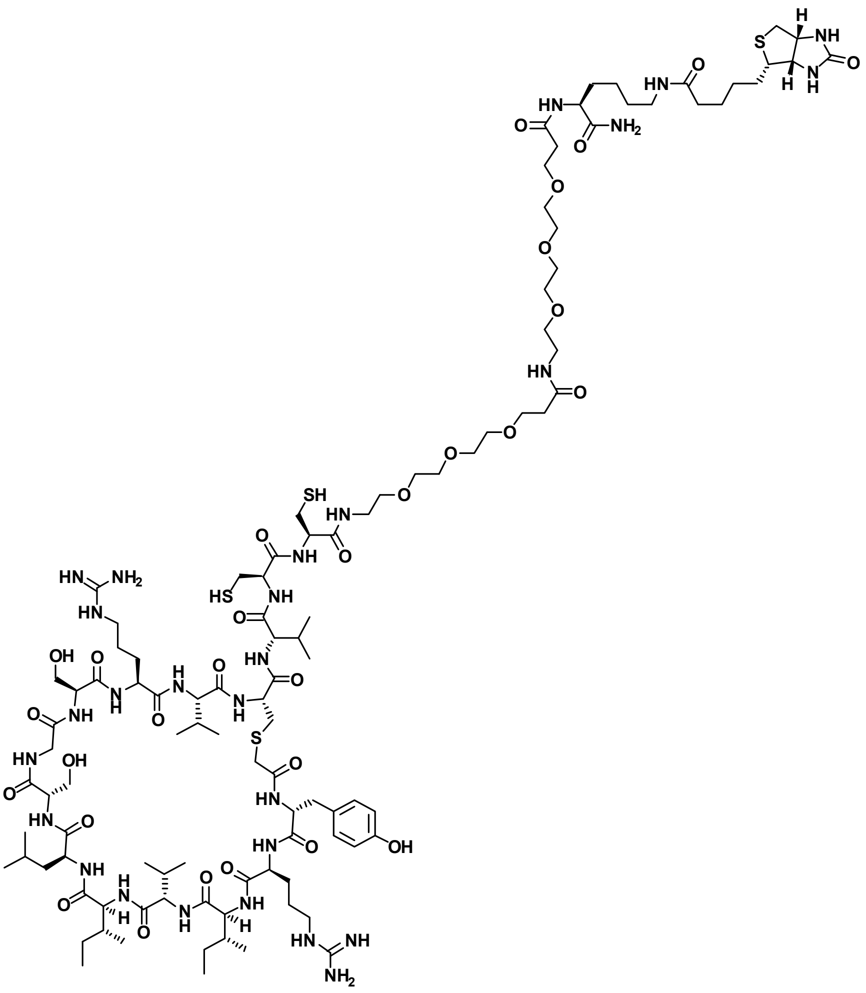

**Biotinylated 5:** Rink amide resin (90 mg, 50 µmol, 0.56 mmol $g^{-1}$) was loaded with Fmoc-Lys(Alloc)-OH according to general procedure **A**. The peptide was then synthesized by iterative Fmoc solid-phase peptide synthesis (Fmoc SPPS) on the Syro I peptide synthesizer according to general procedure **A**, with chloroacetic acid as the final coupling step to generate the resin-bound sequence yRIVILSGSRVCVC(Acm)C(Acm)-[$PEG_3$]-[$PEG_3$]-K(Alloc). Notably, the non-standard Fmoc-protected amino acid, Fmoc-Cys(Acm)-OH, was used as the Fmoc-Xaa-OH for two cysteine residues to enable selective cyclization of the Cys residue closest to the *N*-terminus. A portion (25 µmol) of the peptide was treated with $Pd(PPh_3)_4$ (5 mg, 5 µmol, 0.2 eq.) and $PhSiH_3$ (62 µL, 500, µmol, 20 eq.) in $CH_2Cl_2$ (0.75 mL, 0.03 M with respect to resin loading) and shaken at room temperature for 20 min. The resin was then drained and washed with $CH_2Cl_2$ (10 × 5 mL) and DMF (5 × 5 mL). The resin was then treated with a solution of (+)-biotin *N*-hydroxysuccinimide ester (17 mg, 50 µmol, 2 eq.) and *i*$Pr_2NEt$ (17 µL, 100 µmol, 4 eq.) in DMF (0.25 mL, 0.1 M with respect to resin loading) and shaken at room temperature for 16 h. The resin was then drained and washed with DMF (5 × 5 mL), $CH_2Cl_2$

(5 × 5 mL) and DMF (5 × 5 mL). The resin was then treated with a solution of 5 vol.% $Ac_2O$ and 10 vol.% *i*$Pr_2NEt$ (10 vol.%) in DMF (2.5 mL) for 3 min at room temperature before being washed with DMF (5 × 5 mL), $CH_2Cl_2$ (5 × 5 mL), DMF (5 × 5 mL) and $CH_2Cl_2$ (5 × 5 mL). The peptide was cleaved from resin by general procedure **C** and cyclized according to general procedure **D**. Purification by preparative HPLC (100% $H_2O$ with 0.1 vol.% TFA over 10 min then 0 to 50 vol.% MeCN in $H_2O$ with 0.1 vol.% TFA over 50 min), followed by concentration *in vacuo*, afforded the Acm-protected peptide as a white solid residue. To a solution of the peptide (20 mg, 7.2 µmol) in 1:1 MeCN/$H_2O$ (5 mL) was added silver acetate (60 mg, 0.36 mmol, 50 eq.). The resulting suspension was shaken at room temperature for 1 h, before dithiothreitol (62 mg, 0.40 mmol, 55 eq.) was added. The suspension was centrifuged (5 min, 22 ºC, at 5000 rcf), collecting the supernatant. The pellet was reagitated with DMSO (5 mL) and centrifuged (5 min, 22 ºC, at 5000 rcf), collecting the supernatant. The combined supernatants were purified by preparative HPLC (100% $H_2O$ with 0.1 vol.% TFA over 10 min then 0 to 50 vol.% MeCN in $H_2O$ with 0.1 vol.% TFA over 50 min), followed by lyophilization, afforded the pure peptide as a white fluffy solid (4.2 mg, 6%). **HRMS:** Calculated for $[C_{107}H_{186}N_{28}O_{30}S_4+2H]^{2+}$: 1235.63808, found: 1235.63918. **LRMS:** (+ESI) *m/z* 1236.3 $[M+2H]^{2+}$, 824.5 $[M+3H]^{3+}$, 618.6 $[M+4H]^{4+}$. **Analytical HPLC:** $R_t$ = 15.7 min (1 to 80 vol.% MeCN in $H_2O$ with 0.1 vol.% TFA over 30 min, λ = 214 nm).

A) Analytical HPLC Trace (λ= 214 nm)

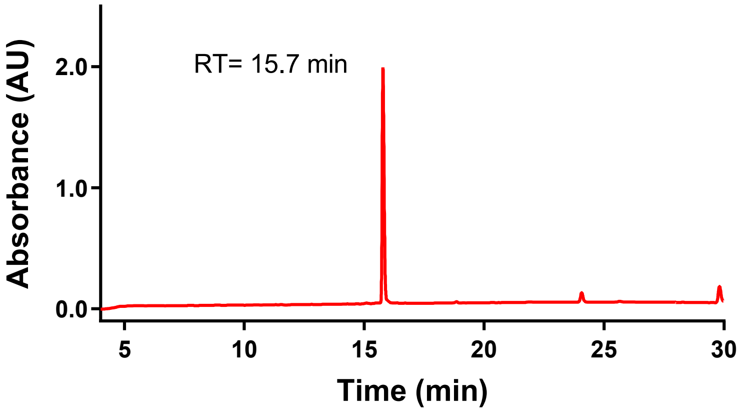


B) ESI MS (+)

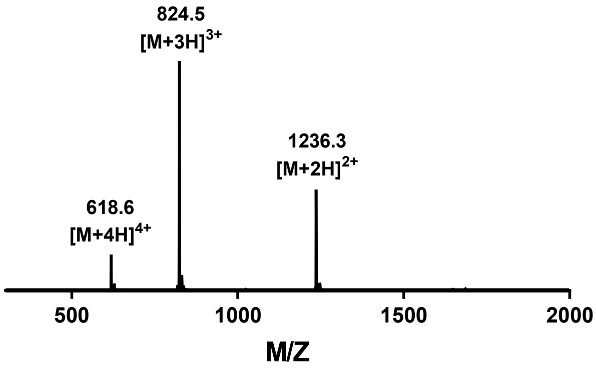

### Mammalian cell expression of SARS-CoV-2 proteins

#### Expression constructs

The expression constructs for the SARS-CoV-2 spike protein (1–1208), the receptor binding domain (RBD) of the spike protein (319–541) used for ELISA assays, the Avitagged™ RBD protein (328–531) used for SPR/BLI assays and the Avitagged™ soluble domain of ACE2 (1–614) were generated as described previously.[3] The expression vectors with soluble trimeric SARS-CoV-2 spike protein and SARS-CoV-2 RBD (319–541) protein were kindly provided by Dr Florian Krammer (Icahn School of Medicine, Mt Sinai, these constructs were used for ELISA-based detection of viral antigens).[4,5] The SARS-CoV-2 spike construct is characterized by the presence of the proteins native signal peptide (residues 1–14) to enable secretion, Pro substitutions at residues 986 and 987, a GSAS substitution at the furin cleavage site (682–685) and a *C*-terminal $His_6$-tag to allow affinity purification. The SARS-CoV-2 RBD (319–541) construct possesses the spike proteins native signal peptide at the *N*-terminus and a *C*-terminal $His_6$-tag to allow affinity purification. The Avitagged™ RBD protein and ACE2 receptor were cloned into the pCAGGS and pcDNA3.1 expression vectors, respectively. Both proteins were *C*-terminally tagged with a $His_9$-tag and an Avitag™ to enable purification and biotinylation, respectively. The Avitagged™ RBD protein was cloned with and *N*-terminal IgK leader sequence and the Avitagged™ ACE2 construct included the proteins native signal peptide (residues 1–18) to allow secretion of the proteins upon expression. The expression construct for *E. coli* biotin ligase, BirA, was cloned with an *N*-terminal Cd4 signal peptide into the pcDNA3.1 vector to allow for enzymatic biotinylation of the Avitag™ when co-transfected with the Avitagged™ constructs.

#### Expression and purification

All proteins were expressed in suspension-adapted Expi293F™ cells using 25 kDa linear polyethyleneimine (PEI). Expi293F™ cells were grown to a density of $3.0 \times 10^6$ cells $mL^{-1}$ in their specified medium and transfected with pre-formed DNA:PEI complexes (2 μg $mL^{-1}$ DNA and 8 μg $mL^{-1}$ PEI). The SARS-CoV-2 RBD and ACE2 receptor transfected cells were incubated for 72 h at 37 °C, while the SARS-CoV-2 spike transfected cells were incubated for 24 h at 37 °C and then transferred to 32 °C for a further 72 h, with 5% $CO_2$ and horizontal orbital shaking at 130 rpm. Medium, containing the secreted proteins, was harvested by centrifugation at 4000 g for 20 min. Upon centrifugation, supernatants were supplemented with

20 mM HEPES pH 8.0 and subject to immobilized Ni-affinity chromatography using Ni-NTA agarose equilibrated in 20 mM $NaH_2PO_4$ pH 8.0, 500 mM NaCl and 20 mM imidazole. Proteins were eluted using a buffer containing 20 mM $NaH_2PO_4$ pH 7.4, 300 mM NaCl, and 500 mM imidazole. Protein-containing fractions were pooled and concentrated to a small volume prior to loading on a Superdex 200 10/30 GL column (GE Healthcare). Proteins were eluted using 20 mM HEPES pH 7.4 and 150 mM NaCl. Protein purification was analysed by SDS-PAGE and multiple angle laser light scattering (MALLS).

For biolayer interferometry (BLI) and surface plasmon resonance (SPR), ACE2 and SARS-CoV-2 RBD were enzymatically biotinylated at the *C*-terminal Avitag™ by co-transfecting the proteins with the BirA expression construct and supplementing the culture media with 100 µM biotin during expression.

The SARS-CoV-2 RBD (319–541) protein used for the RaPID selections was chemically biotinylated at the *N*-terminus using EZ-link™ NHS-Biotin by performing the reaction in a buffer comprising 10 mM HEPES pH 6.5 and 150 mM NaCl at 4 °C overnight. The EZ-link™ NHS-Biotin was added at a 10-molar excess to the SARS-CoV-2 RBD protein.

#### RBD Protein - SPR Studies

SPR measurements were performed using a Biacore™ T200 (GE Healthcare) with 10 mM HEPES pH 7.4, 150 mM NaCl, 0.005% (w/v) Tween® 20 and 0.1% DMSO as running buffer. The sensorchip surface of a CM5 chip (GE Healthcare) was activated using NHS/EDC before immobilization of SARS-CoV-2 RBD or SARS-CoV-2 spike protein in acetate buffer pH 4.0. Alternatively, streptavidin was immobilized on a CM5 chip in acetate buffer pH 4.5 before capture of biotinylated SARS-CoV-2 RBD protein. All experiments were conducted at 25 °C in single cycle kinetics mode with a flow rate of 50 µL $min^{-1}$. Data was analysed using the Biacore™ Insight Evaluation Software.

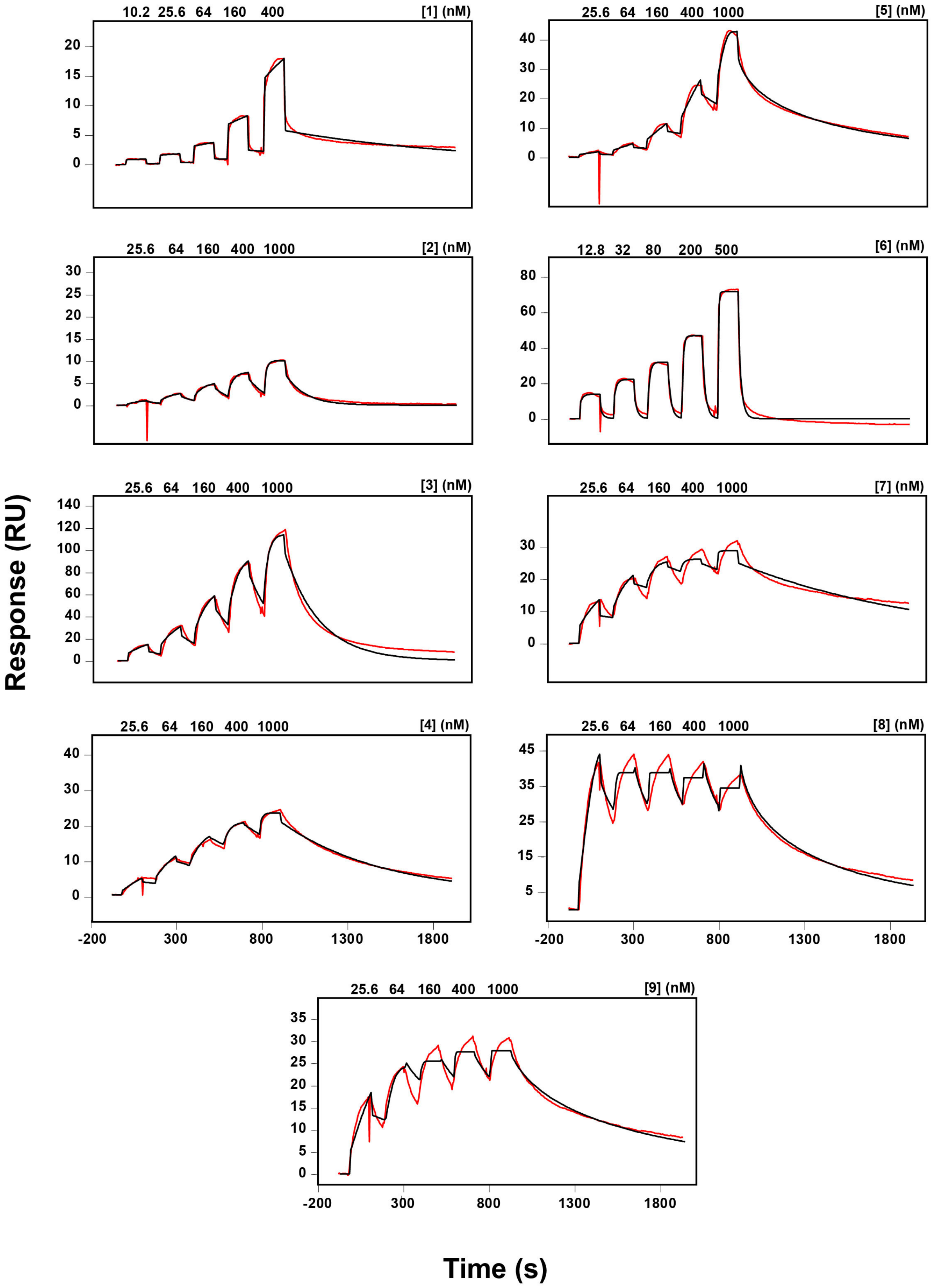


**Figure S1**. Representative SPR sensorgrams for binding of peptides **1–9** to the SARS-CoV-2 RBD protein. Fits to a 1:1 binding model (black) are shown for each sensorgram.

#### Competition Biolayer Interferometry (BLI)

All assays were performed using a BLItz system (ForteBio) at room temperature using 10 mM HEPES pH 7.4, 150 mM NaCl, 0.005% (w/v) Tween® 20 and 0.1% DMSO as kinetic buffer. Biotinylated ACE2 protein was immobilized onto a streptavidin Dip and Read™ biosensor (ForteBio) for 300 s. The sensor was subsequently dipped into kinetic buffer for 30 s followed by a dip into solution containing SARS-CoV-2 RBD (100 nM) and peptide (at concentrations of 0, 1.25 µM, 2.5 µM, 5 µM and 10 µM) for 120 s. To allow for dissociation, the biosensor was dipped into kinetic buffer for an additional 120 s. Data were analysed using the ForteBio Data analysis software and processed by aligning the beginning of the association and dissociation.

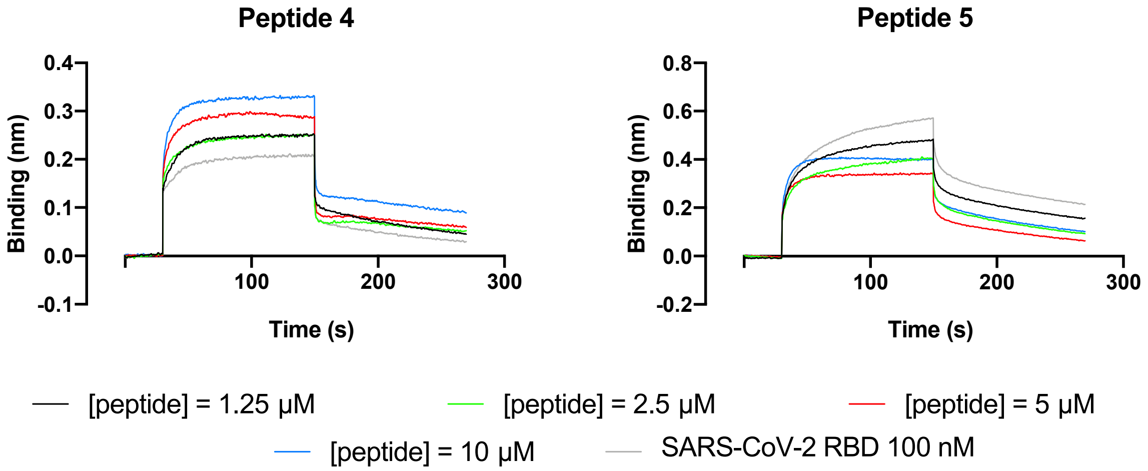


**Figure S2**. Representative BLI sensorgrams for competition of peptides **4** and **5** with ACE2 for binding to SARS-CoV-2 RBD protein. Biotinylated ACE2 protein was immobilized on BLI tips and dipped into a protein solution containing a constant concentration of SARS-CoV-2 RBD (100 nM) and a varying concentration of peptide (1.25 µM, 2.5 µM, 5 µM and 10 µM).

#### Spike Protein - SPR Studies

SPR measurements were performed using a Biacore™ T200 (GE Healthcare) with 10 mM HEPES pH 7.4, 150 mM NaCl, 0.005% (w/v) Tween® 20 and 0.1% DMSO as running buffer. The sensorchip surface of a CM5 chip (GE Healthcare) was activated using NHS/EDC before immobilisation of SARS-CoV-2 spike protein in acetate buffer pH 4.0. All experiments were conducted at 25 °C in single cycle kinetics mode with a flow rate of 50 µL min$^{-1}$. Data was analysed using the Biacore™ Insight Evaluation Software.

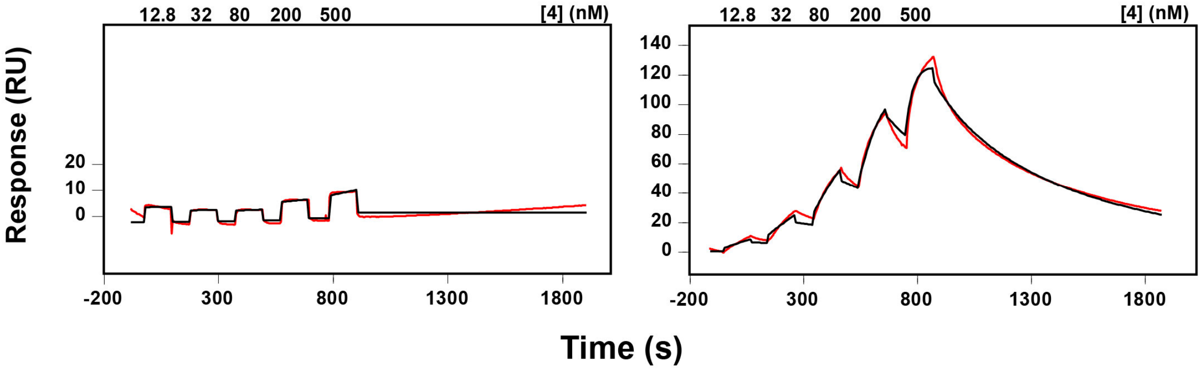


**Figure S3**. Representative SPR sensorgrams for binding of peptide **4** to the SARS-CoV-2 spike protein (left) and the SARS-CoV-2 RBD protein (right). Fits to a 1:1 binding model (black) are shown for each sensorgram.

### Viral Neutralization Assay

##### Cell line

Vero-E6 (ATCC® CRL-1586™) were maintained in Minimal Essential Media (Invitrogen) supplemented with 10% FBS and subcultured according to the supplier's instructions.

##### SARS-CoV-2 virus inhibition assay

A high content fluorescence microscopy approach was used to assess the ability of various RBD cyclic peptide ligands to inhibit SARS-CoV-2 infection in permissive cells. Inhibitors were initially diluted in cell culture medium (MEM-2% FCS) to make 2 × working stock solutions and then serially diluted further in the above media to achieve a 2-fold dilution series. 50 µL of each dilution was mixed with an equal volume of virus solution at $8 \times 10^3$ TCID50/mL and incubated for one hour at 37 °C. Meanwhile, VeroE6 cells were trypsinized and plated in 384-well plates (Corning #CLS3985) at $5 \times 10^3$ cells per well in 40 µL media. After 1 hour, 40 µL of virus-antagonist mix was added to the cells for a final well volume of 80 µL.[6] Plates were incubated at 37 °C, 5% $CO_2$ for a further 72 hours following which the cells were stained with NucBlue dye (Invitrogen, #R37605) according to manufacturer's instructions. Stained cells were then imaged with InCell high throughput microscope (Cytiva) and cell numbers enumerated using InCarta high content image analysis software (Cytiva) to give a quantitative measure for cytopathic effect. Virus inhibition/neutralization was calculated as %N= (D-(1-Q)) ×100/D, where; "Q" is the value of nuclei in test well divided by the average number of nuclei in untreated uninfected controls, and "D"=1-Q for the wells infected with virus but untreated with inhibitors. Thus, the average nuclear counts for the infected and uninfected cell controls get defined as 0% and 100% neutralization respectively. To account for cell death due to drug toxicity cells treated with inhibitor alone and without virus were included in each assay. The % neutralization for each drug concentration in infected wells was normalized to % neutralization in wells with equivalent amount of drug but without the virus to yield the final neutralization values for each condition.

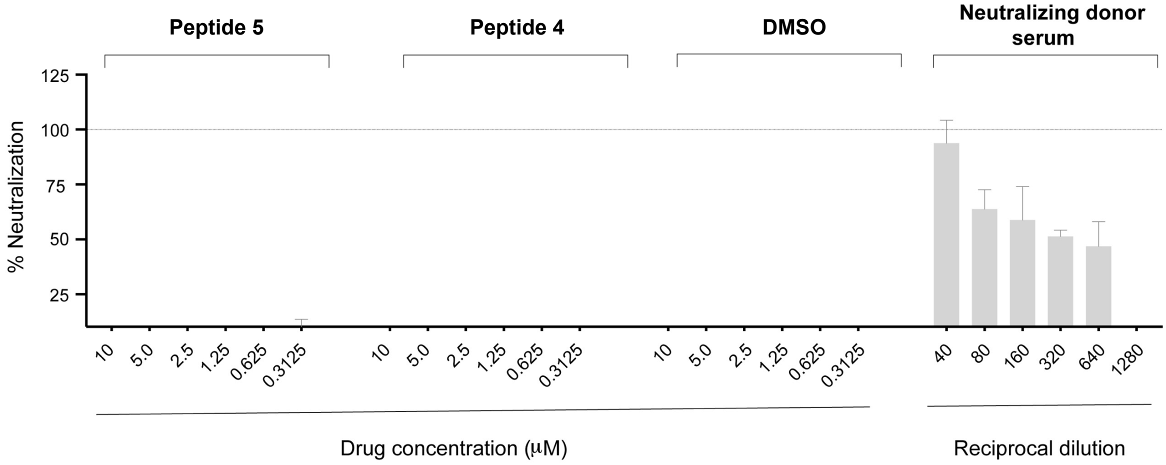


**Figure S4**. SARS-CoV-2 neutralization assay. Varying concentrations of RBD inhibitors were incubated with SARS-CoV-2 and added to Vero E6 cells. Virus cytopathic effect was quantified at 72 hours post infection. Convalescent donor serum served as a positive control for neutralization while DMSO treatment was used as a negative control.

### Pseudovirus Neutralization Assay

##### Expression constructs and cells

The SARS-CoV-2 spike open reading frame, containing an 18 amino acid C-terminal truncation cloned into pCG1 was kindly provided by Prof Stefan Pohlmann.[7] The GFP-luciferase vector construct used was described in Tiffen *et al.*,[8] and lentivirus packaging and helper constructs used were described in Koldej *et al.*[9] 293T cells (CRL-3216) were maintained in Dulbecco's modified Eagle medium (DMEM) and ACE2 overexpressing 293T cells were generated by transducing cells with lentivirus expressing hACE2. The hACE2 ORF was cloned into a 3rd generation lentiviral expression vector, pRRLsinPPT.CMV.GFP.WPRE[10] to create a novel expression plasmid PPT-ACE2. The plasmid was co transfected into HEK293T cells along with psPAX2 (NIH AIDS repository) and pMD2.G (Addgene plasmid # 12259) using polyethylenimine as previously described.[11] Supernatants were collected 72 hours post transfection and transduced into HEK293T cells to generate stable ACE2-HEK293T cell line which was then single cell sorted to generate clonal populations.

**Pseudovirus neutralisation assay**

Replication-deficient pseudotyped lentivirus particles were generated by co-transfecting the GFP-luciferase vector and SARS-CoV-2 expression construct with lentivirus packaging and helper constructs into 293T cells using Fugene HD (Promega) as previously described.[12] Pseudovirus particles were then incubated with RBD peptide binders at various concentrations (50 μM to 0.19 nM) at 37 °C for 1 h prior to spinoculation (800 rcf) of ACE2 over-expressing 293T cells. 72 h post-transduction, cells were fixed and stained with Hoechst 33342 (NucBlue™ Live ReadyProbes™ Reagent, Invitrogen) as per the manufacturer's instructions, imaged used an Opera Phenix high content screening system (Perkin Elmer) and the percentage of GFP positive cells was enumerated (Harmony® high-content analysis software, Perkin Elmer). BD-218, a human IgG1 monoclonal antibody identified by high-throughput single-cell sequencing of a SARS-CoV-2 convalescent patient's B cells by Cao *et al.* was used as a positive control for neutralisation.[13]

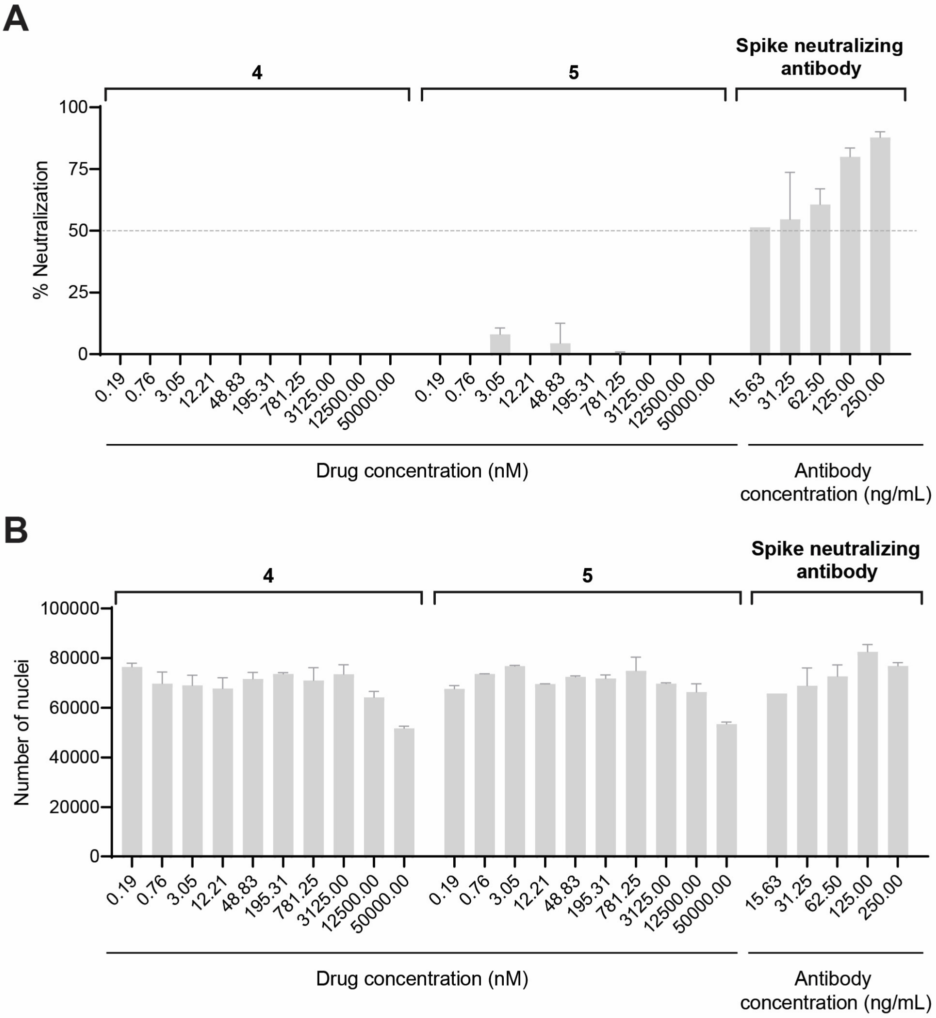


**Figure S5.** SARS-CoV-2 pseudovirus neutralization assay. Varying concentrations of RBD inhibitors were incubated with pseudovirus particles expressing the truncated SARS-CoV-2 spike then added to ACE2 expressing 293T cells. **A.** The GFP reporter plasmid contained in the SARS-CoV-2 pseudovirus particles allowed enumeration of transduced cells 72 h post-transduction. **B.** Cytotoxicity was evaluated by enumeration of nuclei stained with Hoechst 33342. The anti-SARS-CoV-2 spike monoclonal antibody BD-218 was included as a positive control for neutralization.

### Quantitative detection of SARS-CoV-2 Spike and RBD protein by ELISA

Detection of the spike trimer ectodomain and the spike RBD protein from SARS-CoV-2 was performed by either capture or direct ELISA. For antigen capture ELISA, high-binding ELISA plates (Corning Falcon) were coated with monoclonal antibody against SARS-CoV-2 spike protein, BD-218[13] (1 μg/mL), in carbonate/bicarbonate coating buffer (0.05M, pH 9.6) overnight at 4 ºC. After washing with PBS (POCD) with 0.05% v/v/ Tween20 (Sigma), plates were blocked with 1% w/v BSA (Bovogen) in PBS (1 h, 37 ºC). Spike or RBD protein (at a range of dilutions) in PBS was captured (1 h, 37 ºC), the plate washed as before, and antigen detected with biotinylated **4** or **5** (1 µg/mL in PBS, 1 h, 25 ºC). Following washing and labelling of bound biotinylated peptides with streptavidin-horseradish peroxidase (in 1% BSA/PBS, 20 min, 25 ºC), washed ELISA plates were developed with tetramethylbenzidine substrate (Sigma). The reaction was neutralized with 2 M HCl, then absorbance read at 450 nm (570 nm reference wavelength) (Tecan Infinite M1000 PRO). For direct ELISA, ELISA plates were coated as above with spike or RBD protein (at a range of dilutions), blocked and detected with biotinylated **4** or **5** as above. In both ELISAs, non-specific binding of **4** and **5** was assessed by determining detection of unrelated protein, bovine serum albumin (BSA; Bovogen). Biotinylated **4** provided quantitative detection of spike protein at a minimum of 31.25 ng/mL by either antigen capture or direct ELISA, whereas sensitivity for RBD detection was less than 15 ng/mL (**Figure S6**).

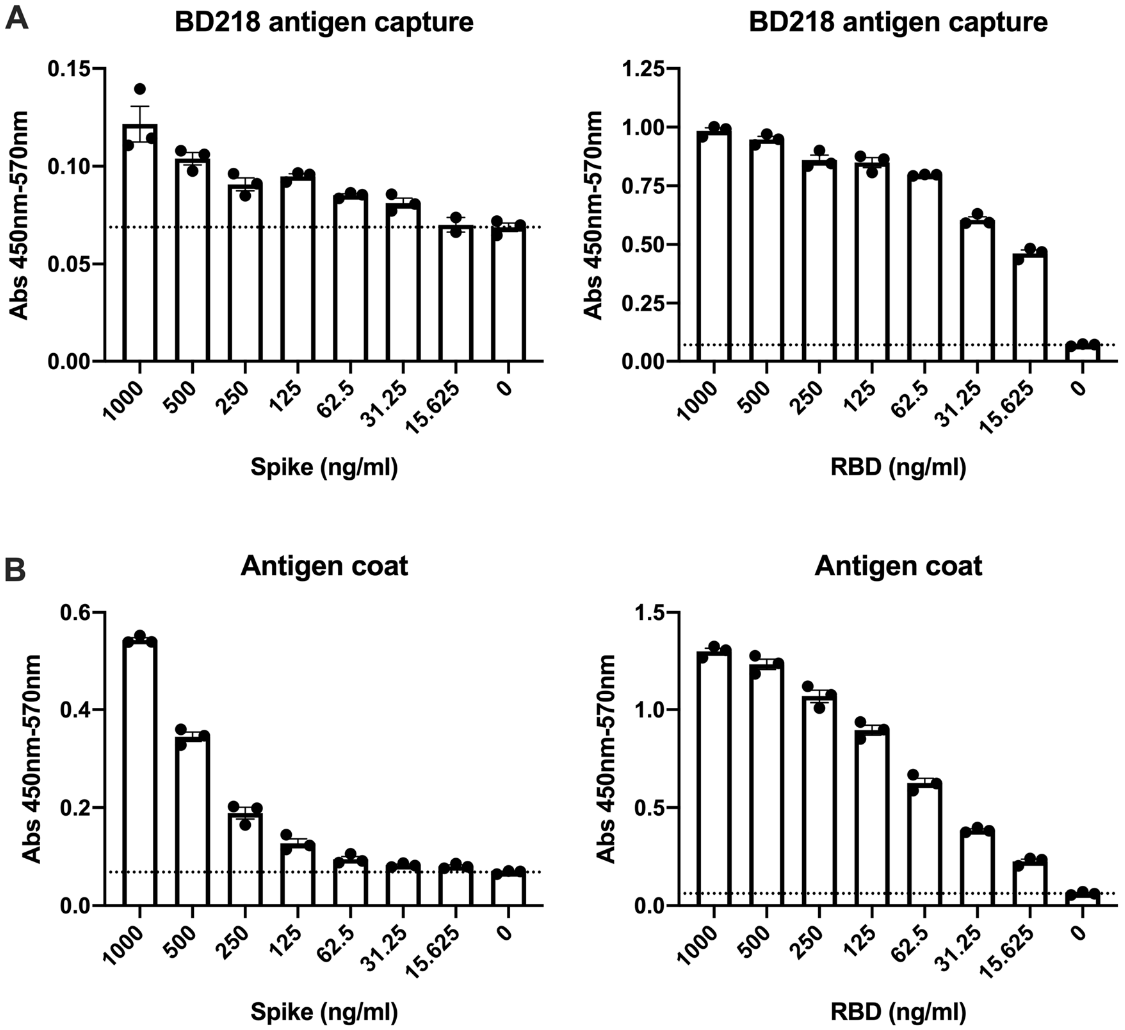


**Figure S6.** Quantitative detection of the SARS-CoV-2 spike protein and RBD by ELISA using biotinylated **4**. **A)** Antigen capture ELISA. Plate was coated with antibody BD-218, blocked with BSA, then either spike or RBD (2-fold dilution series 1000–15.625 ng/mL) was captured and detected with biotinylated **4. B)** Direct ELISA. Plate was coated with either spike or RBD protein (dilution series 1000–15.625 ng/mL), blocked with BSA, and viral antigen detected with biotinylated **4**. Individual data points for technical triplicates are shown, with SEM.

### X-ray Crystallography

Crystals of the SARS-CoV-2 RBD (319–541) protein in complex with cyclic peptide **4** were obtained using hanging drop vapour diffusion. Purified SARS-CoV-2 RBD-$His_6$ in 20 mM HEPES 7.4, 150 mM NaCl were incubated with a 1.2 molar excess of cyclic peptide **4** for 15 minutes at room temperature. Crystals were obtained by mixing 1 μL of SARS-CoV-2 peptide **4** complexes with 1 μL of reservoir solution containing 200 mM ammonium sulfate, 24% PEG

4000, 12% glycerol. Iterative rounds of streak seeding produced diffraction-quality crystals, which were cryoprotected in reservoir solution supplemented with 10% glycerol before flash freezing in liquid nitrogen.

X-ray diffraction data were collected using the MX2[14] beamline at the Australian Synchrotron at 100 K. Data were integrated using XDS[15] and scaled with Aimless.[16] Initial phases were obtained by molecular replacement (Phaser)[17] using SARS-CoV-2 RBD from PDB ID 6M0J (residues 333–516) as the search model. After iterative cycles of manual model building (COOT)[18] and refinement (Phenix),[19] clear difference density was observed near the SARS-CoV-2 RBD *N*- and *C*-termini (Figure S7A) and the cyclic peptide **4** could be modelled in the difference maps (Figure S7B). The final model has good geometry (Table S1) with a MolProbity score[20] of 1.81.

**Table S1**: Data collection and refinement statistics for the crystal structure of RBD in complex with peptide **4**

| | RBD-**4** |
|---|---|
| Spacegroup | H32 |
| Unit cell | |
| Dimensions a,b,c (Å) | 284.92, 284.92, 156.02 |
| Angles α,β,γ (°) | 90, 90, 120 |
| **Data collection** | |
| Wavelength (Å) | 0.9537 |
| Resolution | 48.85-3.96 |
| Observed reflections* | 409593 |
| Unique reflections*# | 21115 (4246) |
| Completeness (%)*# | 99.6 (98.2) |
| Multiplicity*# | 19.4 (19.4) |
| $R_{pim}$*# | 0.075 (0.256) |
| $CC_{1/2}$ *# | 0.996 (0.959) |
| I/σ(I) *# | 7 (1.7) |
| Wilson B (Å$^2$) | 110 |
| **Refinement** | |
| Resolution | 48.85-3.96 |
| $R_{work}/R_{free}$ | 25.7/28.2 |
| Protein molecules/asu | 5 |
| Peptide molecules/asu | 5 |
| Number of atoms | |
| Protein | 7956 |
| Ligand | 98 |
| Peptide | 640 |
| Ramachandran plot | |
| Favored (%)^ | 95.3 |
| Outliers (%)^ | 0.0 |
| B-factors (Å$^2$) | |
| Protein (Chains A,B,C,D,E) | 157, 131, 160, 155, 177 |
| Peptide (Chains R,S,T,U,V) | 150, 138, 182, 190, 210 |
| Root mean square deviation | |
| Bond lengths (Å) | 0.004 |

| | |
|---|---|
| Bond angles (°) | 0.737 |
| PDB ID code | 7L4Z |

*Output from Aimless
#Values in parenthesis are of the highest resolution shell
^Calculated by Molprobity

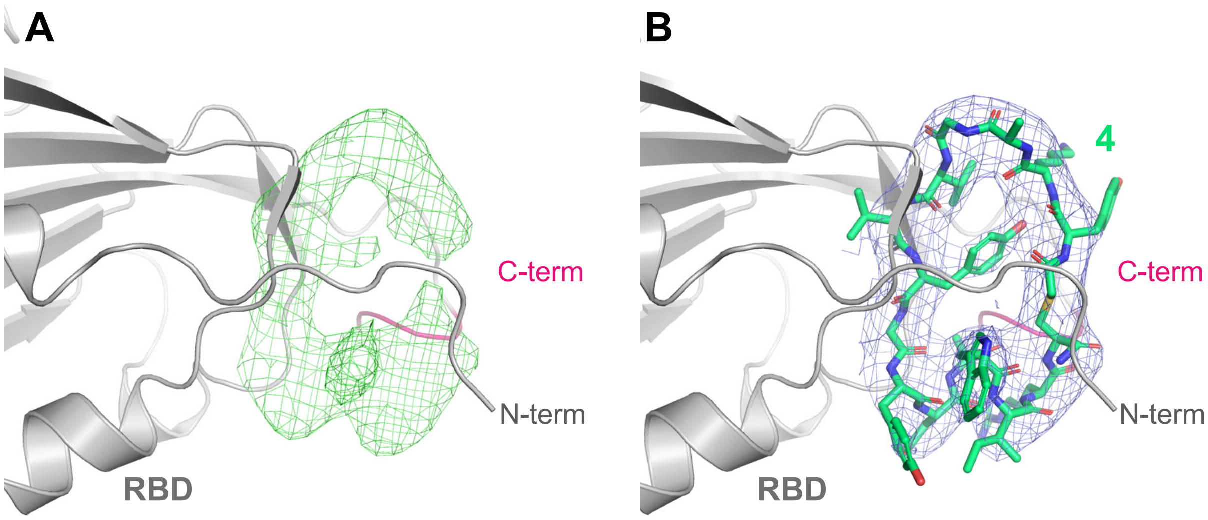


**Figure S7**. (A) Difference $F_o$-$F_c$ density between the N- and C-termini of RBD contoured at 2.5 rmsd (green mesh), after molecular replacement and initial rounds of model building/refinement but before any peptide was built into the maps. RBD is shown in grey cartoon with C-terminal residues $^{523}$TVCG$^{526}$ highlighted in pink. (B) Final 2$F_o$-$F_c$ density contoured at 1 rmsd (blue mesh) with peptide **4** shown as green sticks.
